# A membrane-associated light harvesting model is enabled by functionalized assemblies of gene-doubled TMV proteins

**DOI:** 10.1101/2022.09.08.507180

**Authors:** Jing Dai, Kiera B. Wilhelm, Amanda J. Bischoff, Jose H. Pereira, Michel T. Dedeo, Derek M. García-Almedina, Paul D. Adams, Jay T. Groves, Matthew B. Francis

**Affiliations:** Department of Chemistry, University of California, Berkeley, California 94720, United States; Technology Division, Joint BioEnergy Institute, Emeryville, CA, USA; Molecular Biophysics and Integrated Bioimaging Division, Lawrence Berkeley National Laboratory, Berkeley, CA, USA; Department of Bioengineering, University of California, Berkeley, Berkeley, California 94720, United States

**Keywords:** protein engineering, photosynthesis, supported lipid bilayers, bioconjugation, viral capsid proteins, single molecule imaging

## Abstract

Photosynthetic light harvesting requires efficient energy transfer within dynamic networks of light harvesting complexes embedded within phospholipid membranes. Artificial light harvesting models are valuable tools for understanding the structural features underpinning energy absorption and transfer within chromophore arrays. Most artificial light harvesting complexes are static or in the solution phase, rather than in a two-dimensional fluid environment as in natural photosynthesis. We have developed a method for attaching a protein-based light harvesting model to a supported lipid bilayer (SLB), which provides an extended fluid membrane surface stably associated with a solid substrate. The protein model consisted of the tobacco mosaic viral capsid proteins (TMV) that were gene-doubled to create a tandem dimer (dTMV). Assemblies of dTMV were shown to break the facial symmetry of the double disk to allow for differentiation between the disk faces. Single reactive lysine and cysteine residues were incorporated into opposing surfaces of each monomer of the dTMV assemblies. This allowed for the site-selective attachment of both chromophores for light absorption and a peptide for attachment to the SLB. A cysteine modification strategy using the enzyme tyrosinase was employed for the bioconjugation of a peptide containing a polyhistidine tag for association with SLBs. The dual-modified dTMV complexes showed significant association with SLBs and exhibited mobility on the bilayer. The techniques used herein offer a new method for protein-surface attachment and provide a platform for evaluating excited state energy transfer events in a dynamic, fully synthetic artificial light harvesting system.

**Significance Statement:** Here we have constructed a model photosynthetic membrane containing proteins, chromophores, lipids, and aqueous components, all of which can be modified in their composition. This model is based on an asymmetric disk assembly consisting of engineered tandem dimers of the tobacco mosaic viral capsid protein (dTMV). We have developed methods to achieve dye conjugation and attachment of a supported lipid bilayers (SLB) site selectively on distinct protein surfaces. These dye-labeled protein complexes exhibit mobility on the SLB, resulting in a dynamic model of light harvesting membranes using entirely synthetic components. Additionally, this unique asymmetric assembly of TMV and the facile methods for protein functionalization are expected to expand the tunability of model light harvesting systems.

## Introduction

Photosynthetic light harvesting apparatuses are multi-component machines composed of dynamic networks of light harvesting complexes (1, 2). Energy transfer occurs across networks of multiple dye-bearing membrane protein assemblies working in concert (1, 3). One well-known example of such organization in nature can be found in the purple bacteria *Rhodopseudomonas acidophila*, where light harvesting complex 1 (LH1) is surrounded by and accepts energy from light harvesting complex 2 (LH2) (4). Incoming light is shuttled through LH2 and LH1 before reaching the reaction centers for energy conversion. Numerous artificial light harvesting systems have been developed in pursuit of more efficient photovoltaic and photoelectronic devices and as simplified model systems to probe the underlying structural and chemical features enabling energy transfer in photosynthetic organisms. Substantial effort has gone into creating synthetic light harvesting mimics based on polymers (5), dendrimers (6), nucleic acids (7), and proteins (8) emulating the pigment arrays, orientations, and environmental interactions present within natural light harvesting complexes. These modular, parameterized systems provide insight into the role of pigment–protein interactions (9), the defect tolerance of pigment arrays (10), and quantum entanglement (11), among many other features of energy transfer in photosynthesis (12).

Most of the currently available artificial light harvesting platforms are solution-phase, stand-alone vehicles lacking a lipid component. However, the lipid outer membrane of photosynthetic bacteria and thylakoid membrane of higher organisms, where most photosynthetic light harvesting proteins reside, facilitate energy transfer by allowing reorganization of multi-complex assemblies in response to environmental stimuli. For example, the fluidity of bilayers serves a protective function by allowing light harvesting complexes to aggregate and quench under high light conditions (13). It is also theorized that the major energetic bottleneck of natural light-harvesting processes occurs during inter-assembly electronic energy transfer (EET) (14), which is contingent upon dynamic, non-covalent associations between light harvesting complexes within biological membranes (12). The two-dimensional fluidic movement of light harvesting complexes within lipid bilayers is a missing component in the vast majority of artificial light harvesting systems. The only reported example of a membrane-associated artificial light harvesting model is composed of small molecule chromophores noncovalently embedded in amphiphilic block copolymers (15, 16), and no protein-based systems are available for this purpose. An artificial model composed of membrane-associated, chromophore-bearing protein complexes would closely mimic the fluidity and organization of light harvesting complexes within native organisms. The modularity of this synthetic model would allow us to probe how two-dimensional movement interacts with pigment composition, constraint, and orientation toward elucidating photosynthetic energy transfer mechanisms and pathways.

In this work, we engineer a chromophore-decorated viral capsid protein assembly that exhibits two-dimensional mobility upon association with a lipid bilayer. Synthetic light harvesting systems templated on viral proteins are structurally analogous to naturally occurring complexes due to their composition of self-assembling protein monomers into highly ordered nanoscale structures (8). The tobacco mosaic virus coat protein assembles into helical or circular monomer arrays to which synthetic chromophores can be appended and is highly engineerable, making it a particularly effective light harvesting complex mimic. We have previously used double-disk TMV assemblies to house synthetic individual chromophores and chromophore arrays. The incorporated chromophores demonstrate light absorption and energy transfer (17, 18). Moreover, control over the level of the constraint of chromophores on the TMV surface and within the TMV cavity has shown that the excited state lifetime of these chromophores is modulated by surrounding solvent dynamics and linker compositions (9).

Previously engineered TMV complexes self-assemble from monomer units into *C*_*2*_-symmetric stacks of 2- or 4-disk assemblies, preventing independent functionalization of the top and bottom disks (Fig. 1 a and b). In pursuit of greater facial control over the TMV disk faces, we have constructed a tandem dimer of TMV (dTMV) which breaks the *C*_*2*_ symmetry of previous assemblies and enables orthogonal bioconjugation to the two surfaces of the double-disk assembly, with one bioconjugation handle used for attachment to a chromophore and a second handle used to conjugate the assembly to a fluid supported lipid bilayer (SLB) (19). The SLB provided a two-dimensional surface to which TMV can bind at high density, while preserving free lateral diffusion on the surface to promote energy exchange between disks (Fig. 1c). Moreover, the SLB platform readily enables imaging by standard microscopy (20), a variety of single-molecule imaging (21) and correlation spectroscopic methods (22, 23), as well as advanced spectroscopy (24) techniques to probe the behavior of the membrane-associated, chromophore-labeled TMV assemblies. The SLB platform also facilitates integration with metallic, or other electronic material structures fabricated onto the underlying substrate (25–28).

**Figure 1.**
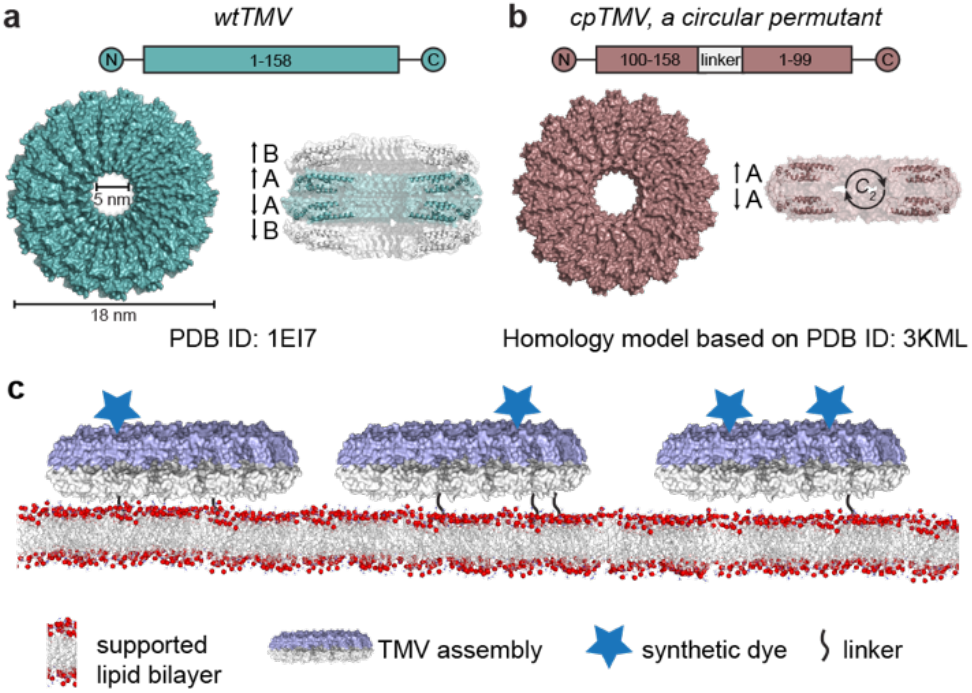
Designing a model system for mobile photosynthetic complexes on a lipid bilayer. (a) The *C*_*2*_-symmetric, four-layer aggregate of wtTMV is composed of layers of 17-monomer disks. The A–A disk pair is *C*_*2*_-symmetric, while the A–B disk pair exhibits translational symmetry with slight structural variation between rings A and B. (b) A circular permutant of TMV, cpTMV, preferentially forms *C*_*2*_-symmetric, two-disk stacks under physiologically relevant conditions. (c) The complex process of photosynthetic energy transfer may be modeled via the covalent attachment of synthetic LHCs to a supported lipid bilayer. The use of synthetic dyes and a synthetic protein complex would allow for stability and tunability of the model system.

## Results

### dTMV was created by a genetic fusion of two rTMV monomers

A TMV coat protein tandem dimer (dTMV) was created by fusing two recombinant TMV (rTMV) monomers on the genetic level (Fig. 2 a–c), with the goal of connecting monomers in the top and bottom disks. Even though circular permutant TMV (cpTMV) forms a more stable double-disk assembly (29), we chose to use rTMV as the dTMV template because the termini are in close proximity to one another on the outer edge of the disks, facilitating their connection by a short linker. In contrast, cpTMV, due to its gene rearrangement, has its two termini in the central pore region where additional linkers could be envisaged to interfere with assembly formation.

**Figure 2.**
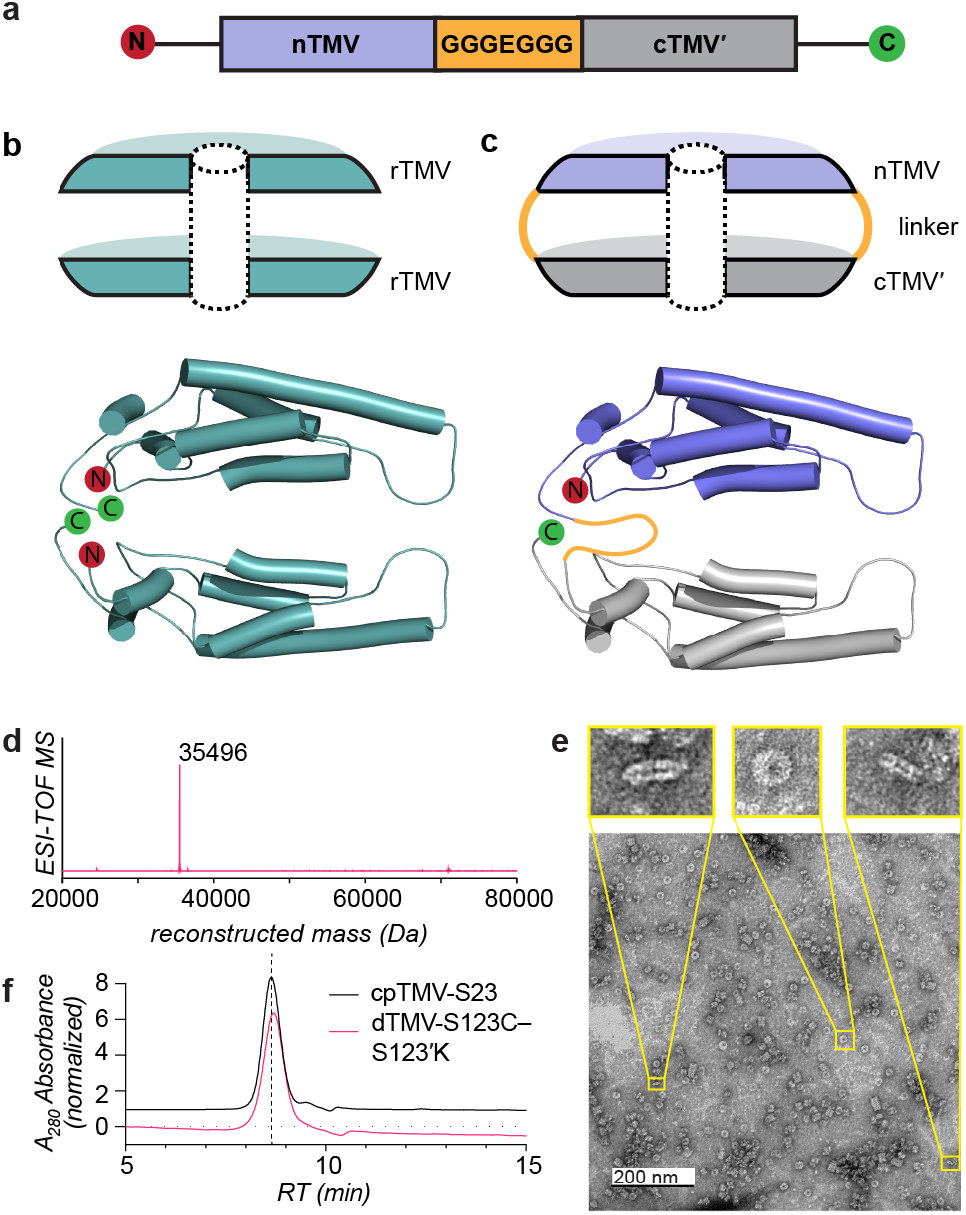
Design considerations and general assembly state of dTMV. (a) The gene map of two fused recombinant TMV (rTMV) monomers contains a flexible linker with the sequence GGGEGGG between the first (nTMV) and second (cTMV′) monomers. (b) A side view of a two-layer disk of wtTMV as well as top and bottom monomers demonstrates that the N- and C-termini of wtTMV are in close proximity to one another on the exterior of the disk assembly. (c) A side view the proposed two-layer disk and tandem dimer of TMV, dTMV, illustrates the linker between the N-terminus of one monomer and the C-terminus of an apposed monomer. (d) The mass of a purified dTMV-S123C–S123′K dimer (fused monomer) matches the expected mass of 35496 Da. (e) TEM images of dTMV-S123C–S123′K reveal a circular face of assembled capsids and stacks of two or multiples of two disks. (f) The assembly state of dTMV-S123C–S123′K was assessed by size exclusion chromatography which showed a size similar to the reported double disk structures of previous TMV constructs with a retention time of 8.5 min.

An experimental evaluation of preliminary linking strategies yielded a well-expressed dTMV construct with a flexible 7-amino-acid linker, GGGEGGG (Fig. S1 a–d). The glutamate residue was included to compensate for the loss of the negative charge of the C-terminus. The two monomers in the tandem dimer, with the N-terminal unit labeled nTMV and the C-terminal unit labeled cTMV′, have the same protein sequence encoded by different nucleotide sequences to ease genetic manipulation. Expression of dTMV resulted in peptide chains of the expected molecular weight (Fig. 2 d). These dTMV monomers self-assembled into intact double-layered disks at the time of expression in *E. coli* (Fig. 2 d and e), as characterized by size exclusion chromatography (SEC) and transmission electron microscopy (TEM).

### dTMV forms stable double disks with each dimer spanning both disks

A crystal structure of the dTMV assembly was obtained and refined to 2.8 Å resolution (Fig. 3). The overall dimension and shape of the dTMV double-disk assembly were similar to those of the AA ring pairs found in both rTMV and cpTMV, with the two disks facing opposite one another (Fig. 3 a and b). As expected, each assembly consisted of 17 dTMV subunits, the same number as found in other TMV disk assemblies (Fig. 3 c). The solved assembly structure strongly implied that the N- and C-terminal TMV subunits occupy positions in separate disks. Though the linkers between the fused monomers were not resolved in the structure, they were not sufficiently long to connect monomers sitting adjacently in the same disk, but could readily connect the C- and N-termini of stacked rTMV units (Fig. S2). However, unlike rTMV and cpTMV, the dTMV assembly was not perfectly *C*_*2*_-symmetric. One disk was flat, similar to the equivalent portion of cpTMV and rTMV (colored purple in Fig. 3 a and b), but the other disk was concave (colored gray in Fig. 3 a and b). The C-terminal segment (residues 77–158) of each rTMV subunit was exposed on the outer surfaces of both the flat and the concave disks, verifying that the S123 sites located in the C-terminal regions and previously used for bioconjugation in rTMV would still be available for modification with fluorophores and bilayer surfaces (17).

**Figure 3.**
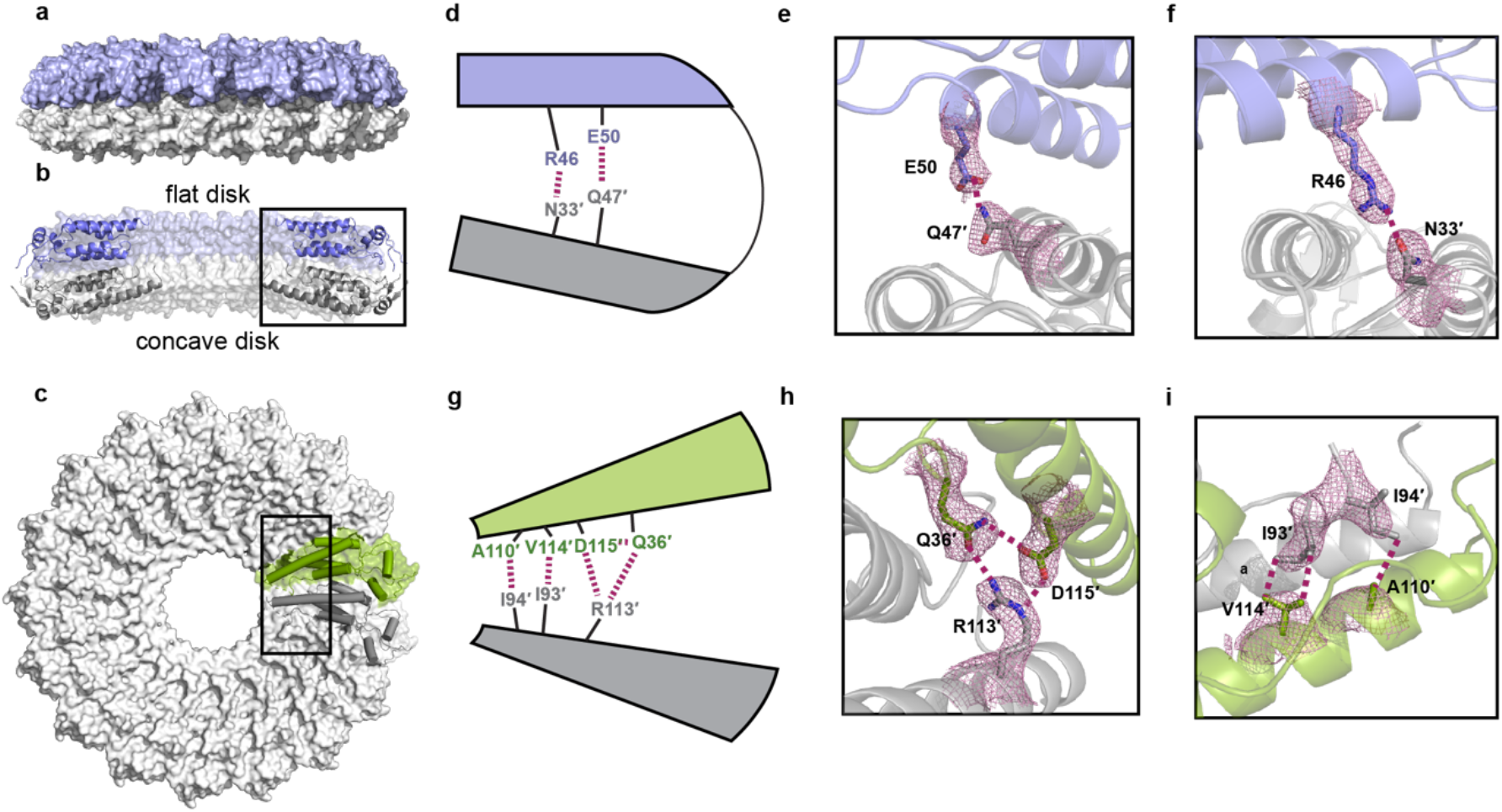
Structure of the dTMV double-disk assembly. Based on the crystal structure, dTMV forms an AA ring pair of the double-disk assembly containing a total of 17 subunits. The side view of the assembly is shown in (a). No orange region for the linker is shown because this region was not well-defined in the crystal structure. (b) The cross-section view reveals that the two disks are not perfectly *C*_*2*_ symmetric as in cpTMV and wtTMV, but instead contain one disk puckering inward (colored grey) relative to the other (colored purple) as determined by x-ray crystallography. The monomers in the concave disk have a more ordered structure in the low-radius region. (c) The top view of the assembly is shown, with a single monomer colored in green. (d) The inter-chain interactions between the flat and concave disks are illustrated. The hydrogen bonds between (e) Glu50 and Gln47 as well as (f) Arg46 and Asn33 are shown together with the 2mF_O_-DF_C_ map contoured at the 1.2σ level (shown as mesh line). (g) The inter-chain interactions in the low-radius regions of two adjacent monomers are illustrated. (h) The hydrogen bonding network among Gln36, Asp115 and Arg113 and (i) the hydrophobic interactions between Val114 and Ile93, as well as Ala110 and Ile94 are shown together with the 2mF_O_-DF_C_ map contoured at the 1.2σ level (shown as mesh line). The narrower intercavity space of dTMV results in direct protein-protein interaction between flat and concave monomers, which is not present in wtTMV or cpTMV.

The lack of resolution in the linker regions limits further identification of nTMV and cTMV′ within the two inequivalent disks. Because both fused monomers in this dTMV construct have the same sequence, the linker stretching from the C-terminus of nTMV to the N-terminus of cTMV′ is the only way to distinguish the two halves of the dimer. However, the C-terminal region of both nTMV and cTMV′ appeared disordered, and though the N-terminal regions are more ordered, both the flat and concave monomers showed positive density extending from their N-terminal regions in the difference electron density map (Fig. S3a). A polder map (30) did not provide any more insight into the linker positions (Fig. S3b). Moreover, tandem dimers within assemblies are not necessarily all oriented in the same direction. They could all be uniformly oriented, with nTMV on the same face and cTMV′ on the opposite face (Fig. S4a), or with nTMV and cTMV′ oriented randomly (Fig. S4b). Either a scrambled orientation or high linker flexibility could account for averaged electron density extending from the N-terminal region of both tandem dimers. Although we cannot conclusively assign nTMV and cTMV′ to the flat or concave disks, for the purpose of clarity in this report we will refer to the residues of the flat monomer as the 1-158 series and those of the concave monomer as the 1′-158′ series.

The crystal structure helps explain the high stability of the dTMV double-disk assembly. The double-disk stayed intact even at pH 9, and a 12 M urea solution was required for disassembly. This is notably higher than the typical 6 M solution used for disassembling cpTMV or rTMV (Fig. S5). We hypothesize that the linkers locking the flat and concave monomers together significantly contribute to stability. In addition, direct protein-protein interactions were observed in the contact regions between the two disks of dTMV assemblies (Fig. 3d). In contrast, rTMV and cpTMV only exhibited weak protein-water-protein interactions in their internal cavity regions (31, 32). The overlaid cross-sections of the dTMV assembly (colored purple and gray in Fig. S6a) and rTMV AA ring pairs (colored orange in Fig. S6a) clearly showed a reduction in the interdisk distance, involving hydrogen bond formation between Glu50 of the flat monomer and Gln47′ of the concave monomer (Fig. 3e), as well as between Arg46 of the flat monomer and Asn33′ of the concave monomer (Fig. 3f).

A superposition of the backbones of the flat and concave monomers suggests that the secondary structures are similar, except for regions close to the central pore (low-radius regions, red box in Fig. S6b). This more organized low-radius region, evidenced by a well-defined electron density, is stabilized by the interaction network (Fig. 3g) only present in the concave disks. The unique interactions between two adjacent concave monomers include a hydrogen bonding network among Gln36′, Asp115′, and Arg113′ (Fig. 3h), hydrophobic interactions between Val114′ and Ile93′, and hydrophobic interactions between Ala110′ and Ile94′ (Fig. 3i).

### Asymmetric dual-functionalization of dTMV allows dye-labeled double disks to conjugate to supported lipid bilayers

To begin the construction of a bilayer-associated dTMV model, mutations were selectively introduced into nTMV and cTMV′ to enable two complementary conjugation reactions. S123 was chosen as the bioconjugation site in both nTMV and cTMV′ because it is surface-exposed in both the flat and concave disks and is the site of previous successful chromophore conjugation (17). Two constructs were created: (1) dTMV-S123C–S123′K, in which the cysteine was introduced in the N-terminal nTMV monomer domain and the lysine was introduced in the C-terminal cTMV′ monomer domain; and (2) dTMV-S123K–S123′C, with the opposite arrangement (Fig. 4a). Aside from these engineered residues, no additional reactive cysteine or lysine residues were expected to be present on the dTMV surface. Both mutants expressed well, producing disks of the expected molecular weight (Fig. 2d) and forming assemblies that were analogous to those of the original dTMV sequence. The double disk morphology was confirmed by SEC in comparison to cpTMV, which is known to form double disks(33) (Fig. 2e), and by TEM (Fig. 2f). The TEM images revealed double disks of the expected diameter (18 nm), and with the characteristic central pore of TMV-based assemblies. Some short, non-helical stacks of disks were also observed in the images, which is consistent with other TMV assemblies (34).

**Figure 4.**
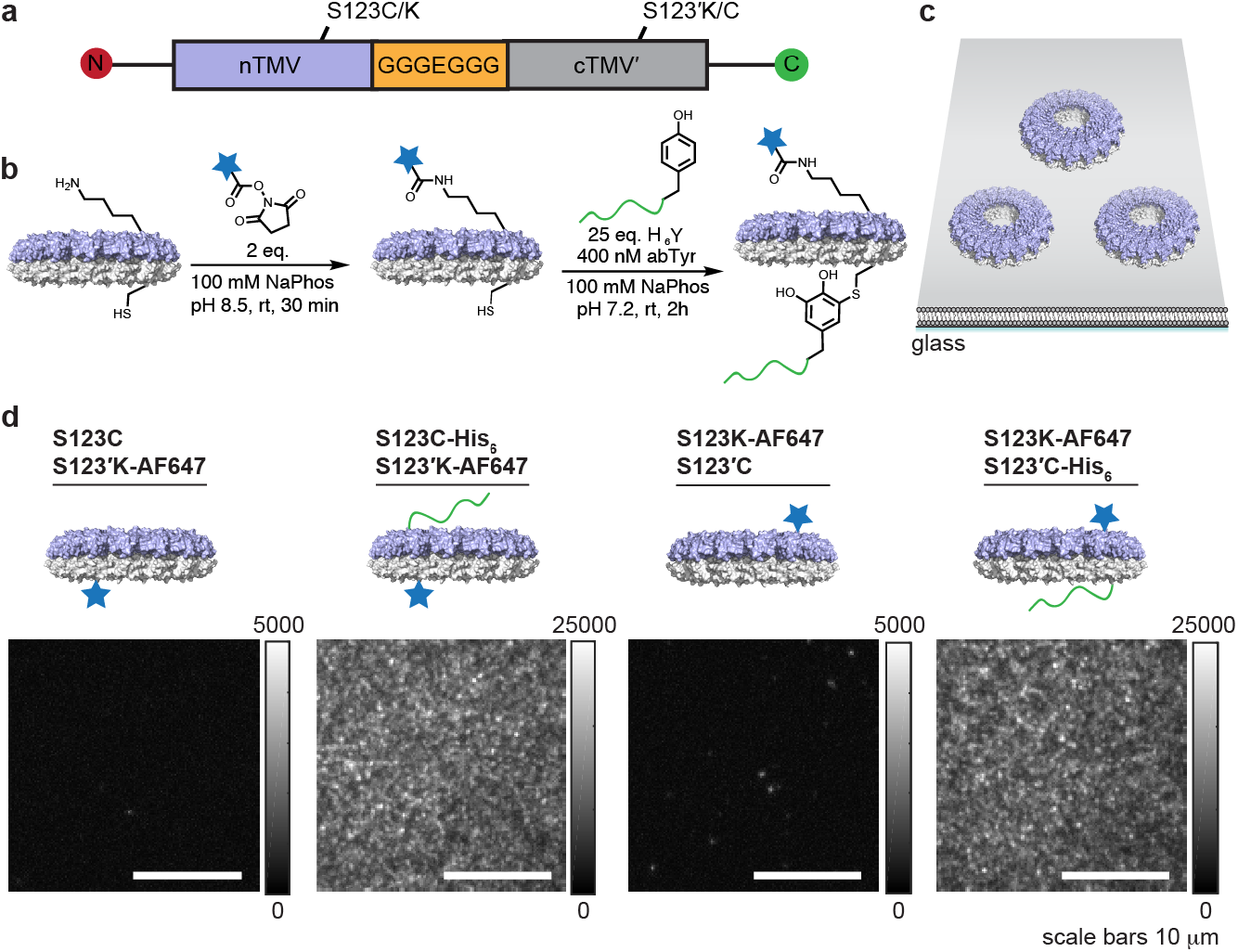
Attachment of dTMV to supported lipid bilayers via a non-expressed His_6_ tag. (a) The amino acid residues lysine and cysteine were installed on each monomer of the dTMV fused dimer via protein engineering. (b) To each engineered dTMV assembly, AlexaFluor 647 (AF647) was attached to the engineered Lys123 or Lys123′ residue using AF647 NHS ester, and the H_6_Y peptide was attached to the Cys123 or Cys123′ residue on the opposite monomer using the enzyme tyrosinase (abTyr). AF647 was used for microscopic visualization, and the H_6_Y peptide was used for attachment to nickel-chelating lipids. (c) A schematic shows dTMV assemblies on the surface of a supported lipid bilayer (SLB). (d) Without the H_6_Y peptide attached to dTMV, neither the dTMV-S123K-AF647–S123′C nor dTMV-S123C–S123′K-AF647 constructs showed significant attachment to bilayers containing Ni-NTA lipids. With similar bioconjugation levels of the H_6_Y peptide (59-60% of monomers modified) and membrane incubation concentration, the AF647-labeled dTMV-S123C-H_6_Y–S123′K-AF647 and dTMV-S123K-AF647– S123′C-H_6_Y assemblies attached to the SLB at similar densities.

Disks composed of uniform constructs were incubated with Alexa Fluor 647 (AF647) *N*-hydroxysuccinimide (NHS) ester to achieve fluorescent labeling of the engineered lysine residue with 5–10 % of monomers labeled (Fig. 4b and Fig. S7a). A construct without the reactive lysine residue, dTMV-S123C-S123′, showed no modification with AF647 under these conditions (Fig. S8a). We then attempted to conjugate the fluorescently-labeled dTMV disks with an exposed cysteine residue to an SLB containing lipids with maleimide head groups. These initial experiments resulted in dTMV sparsely adhering to SLBs, with only marginal increase in density over control bilayers lacking maleimide-functionalized lipids (Fig. S9). Increasing dTMV incubation concentrations > 5-fold did not significantly increase TMV association to the bilayer. Steric hindrance may have prevented the relatively planar surface of the ∼600 kDa dTMV complex from conjugating to the planar SLB at S123(′)C through the desired linkage.

To address this issue, we sought to incorporate a short and flexible His-tag into the dTMV disks to allow their binding to SLBs containing nickel-chelating lipids. However, expression of TMV with an N-terminal His-tag results in altered assembly states and sensitivities to concentration and buffer (32, 35), requiring a different method for adding the His-tag to dTMV. The termini of dTMV are on the periphery of the disks, so the installation of a terminal His-tag could also result in an edge-on orientation of dTMV on the bilayers, rather than the desired face-on orientation. Covalent modification with a His-tag, as opposed to expression, would allow for control over the surface modification site as well as the number of His-tags per 17-monomer assembly.

To modify the existing dTMV constructs at the S123(′)C position with a His-tag, we employed an oxidative coupling reaction using the enzyme tyrosinase for the conjugation of phenol-containing peptides, small molecules, or proteins to cysteine residues on protein surfaces (36–38). Short peptide sequences with a C-terminal tyrosine and an N-terminal His-tag were designed to conjugate to the dTMV double disks at S123(′)C and subsequently coordinate with nickel-nitrilotriacetic acid (Ni^2+^-NTA) lipids in SLBs. Tyrosinase-catalyzed modification of both dTMV-S123C–S123′K and dTMV-S123K–S123′C with a peptide with the sequence HHHHHHY (H_6_Y) resulted in approximately 50% modification, or an average of 8-9 His-tags per double disk assembly, and did not appear to significantly disrupt the assembly state of the double disks (Fig. S7 a–b). Under the same conjugation conditions, no modification with H_6_Y of a construct without the reactive engineered cysteine residue, dTMV-S123K–S123′, was observed (Fig. S8b). This modification proceeded similarly both before and after labeling with the AF647 NHS ester dye. These dTMV-S123(′)K-AF647–S123(′)C-H_6_Y modified double disks specifically conjugated to Ni^2+^-NTA-containing SLBs through multivalent His_6_:Ni^2+^-NTA interactions (Fig. 4 c and d). Densities of 13 and 14 molecules per square micron (μm^-2^) were reached for dTMV-S123C-H_6_Y–S123′K-AF647 and dTMV-S123K-AF647–S123′C-H_6_Y, respectively, with a modest 15 nM incubation for 1 h at room temperature in PBS. The maximum possible density for a single layer of dTMV double disk assemblies on an SLB would be ∼3000 μm^-2^, indicating that the conditions used herein resulted in ∼1/200 of maximum coverage. Higher coverage to facilitate inter-disk energy transfer could be achieved by increasing the dTMV incubation concentration on the SLB (39).

To demonstrate the versatility of this strategy, tyrosinase was also used to modify the surface-exposed cysteine residues of a midsize mammalian protein, bovine serum albumin (BSA) with the H_6_Y peptide after labeling the exposed lysine residues of BSA with AF647 NHS ester. A solution concentration of 100 pM dye-labeled, His-tagged BSA yielded a density of 1.4 μm^-2^ (Fig. S10a–c), which is on par with other proteins containing single His tags (39). This demonstrates the potential versatility of the tyrosinase-catalyzed peptide-peptide coupling strategy and opens the possibility of attaching His-tags and other peptide sequences to proteins with buried N- and C-termini or for which expression with a peptide tag is not feasible or convenient.

### TMV double disks are mobile on supported lipid bilayers

We next used total internal reflection fluorescence (TIRF) microscopy and single molecule analysis to quantitate the mobility and fluorophore labeling of dTMV-S123(′)K-AF647–S123(′)C-H_6_Y double disks on the SLBs. Both constructs – with the His-tag conjugated to the nTMV or cTMV′ units of the dimer – behaved similarly on SLBs. For clarity, results from dTMV-S123C-H_6_Y–S123′K-AF647 are reported in the main text (Fig. 5) and analogous results from dTMV-S123K-AF647–S123′C-H_6_Y are reported in the SI (Fig. S11).

**Figure 5.**
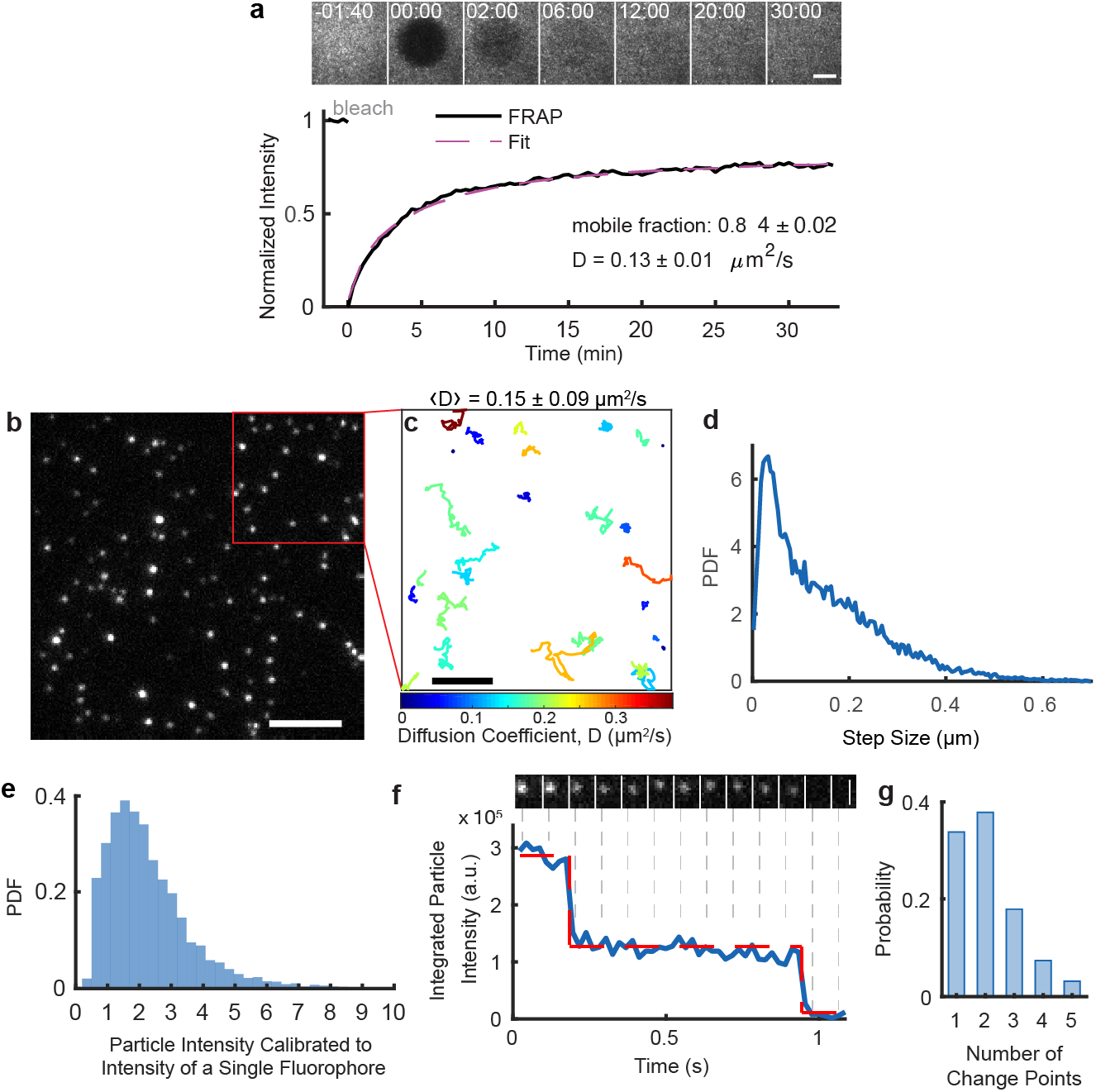
dTMV-S123C-H_6_Y–S123′K-AF647 is mobile on supported lipid bilayers. (a) Fluorescence recovery after photobleaching of a high density (20 μm^-2^) of dTMV on the bilayer indicates that 84% of the dTMV disks on the bilayer are mobile, and that the mobile fraction diffuses at 0.13 ± 0.01 μm^2^/s on average. Scale bar 10 μm. Error denotes 95% CI. (b) A low density (0.2 μm^-2^) of disks on the bilayer enables analysis of single disk particles. Scale bar 5 μm. (c) Tracks of particles depicted in (b) diffusing through time illustrate the varied diffusion rates of particles on the bilayer. Tracks are colored by their diffusion coefficient, D. Scale bar 2 μm. Error denotes standard deviation. (d) The step size distribution of diffusing disks contains contributions from slowly-diffusing particles at smaller step sizes and quickly-diffusing particles at larger step sizes. n = 68,805 steps, n = 2,281 trajectories. (e) The background-subtracted fluorescence intensity distribution of all disks on the bilayer is calibrated to the mean integrated intensity of disks labeled with a single fluorophore. n = 5,934 particles. (f) A representative particle photobleaches in two steps, indicating that two fluorophores are conjugated to that particle. Scale bar 1 μm. (g) The step photobleaching traces of particles that bleach completely over the course of the imaging acquisition are analyzed for the number of change points in their fluorescence intensity. n = 96 particles.

First, fluorescence recovery after photobleaching (FRAP) was used to determine the mobility of dTMV double disks when presented at a high density (14–20 μm^-2^) on a SLB (Movie S1, Movie S2). The fluorescence recovery trace for dTMV-S123C-H_6_Y was normalized and fitted to yield an average diffusion coefficient of 0.13 +/-0.01 μm^2^/s and a mobile fraction of 0.84 +/-0.02 (Fig. 5a) (See Materials and Methods for details). This diffusion coefficient is expected for a large protein complex that coordinates to multiple lipids that diffuse otherwise unhindered in an SLB.(23, 40)

Imaging experiments performed at low double disk density (0.2 μm^-2^) revealed expected heterogeneity in the number of His_6_ tags and dye molecules conjugated to each dTMV double disk (Fig. 5b, Fig. S11b). Tracking of single disk assemblies through time revealed that some particles diffused quickly and some slowly, but the diffusion rates of particles were consistent with time (Fig. 5c, Fig. S11c, Movie S3, Movie S4). This was consistent with double disks being conjugated to His_6_ tags following a broad, Poisson distribution, but the number of contact points between a single disk and the SLB did not vary notably with time. The step size distribution compiled from 2,281 dTMV-S123C-H_6_Y–S123′K-AF647 disks similarly showed contributions from slowly diffusing particles (peak around 0.02 µm step size per 60 ms time interval), moderately diffusing particles (shoulder around 0.2 µm), and quickly diffusing particles (tail extending beyond 0.4 µm) (Fig. 5d).

Analysis of single particle intensities was used to quantitate the number of fluorescent labels on each disk. Background-subtracted particle intensities were normalized to the intensity of a single fluorophore. A histogram of these intensities revealed a labeling efficiency of 2.3 ± 1.5 (SD) fluorophores per dTMV for this sample (Fig. 5f, see Materials and Methods for details). The number of fluorophores conjugated to a specific dTMV double disk particle can be quantitated directly by tracking the fluorescence intensity of the particle through time (Fig. 5g). Fluorophores bleach in discrete steps, so the number of intensity change points until the signal of a particle fully bleaches indicates the number of fluorophores attached to that particle. An analysis of 96 particle intensity traces yielded an average of 2.1 ± 1.0 (SD) fluorophores per dTMV (Fig. 5h), in agreement with the fluorescence intensity distribution.

### Exploring the orientations of the nTMV and cTMV′ domains of dTMV

The fluorescence images of dTMV-S123(′)C-H_6_Y–S123(′)K-AF647 on SLBs (Fig. 4d) did not provide clarity regarding the relative orientations of tandem dimers within dTMV assemblies (Fig. S4 a–b). If similar incubation concentrations of fluorescently labeled dTMV-S123C-H_6_Y–S123′K-AF647 and dTMV–S123K-AF647–S123′C-H_6_Y on Ni-NTA lipid-containing SLBs had resulted in significantly different surface densities, this would have provided evidence that nTMV is always on the flat disk while cTMV′ is always on the concave disk, or vice-versa, due to differing accessibility of the His_6_ tag. However, the observed similarity in surface densities may indicate either that the nTMV and cTMV′ are oriented randomly on the flat and concave disks or that the His-tag is equally accessible to the bilayer on either the flat and concave disk despite structural asymmetry.

To attempt to resolve the relative orientations of the tandem dimers within the dTMV double disk, we labeled dTMV at the S123(′)C site with multiple copies of peptides and proteins. We then imaged the constructs by TEM after applying uranyl acetate negative staining. In the case of a uniform monomer orientation, the attached proteins or peptides would be expected to appear on a single side of the disk, while a scrambled orientation would result in modification of both faces. A longer peptide than H_6_Y, HHHHHHSGGGGY (H_6_SG_4_Y), was used to increase visibility of the peptide by TEM, resulting in 70% modification of both dTMV-S123C–S123′K and dTMV-S123K–S123′C (Fig. S12a). While the results were obscured by the stacking behavior of the dTMV disks, the attached peptide was not clearly resolved in images of either dTMV-S123C-H_6_SG_4_Y–S123′K or dTMV-S123K–S123′C-H_6_SG_4_Y (Fig. S12b). When conjugated to an anti-HER2 nanobody containing a C-terminal SGGGGY tag (nbHER2_Tyr_),(41) dTMV-S123K–S123′C appeared to show examples of nbHER2_Tyr_ conjugation to both sides of the double disk (Fig. S12 c and d). The TEM images as well as fluorescence images of modified dTMV are inconclusive, but suggestive of a random orientation of nTMV and cTMV′ on the flat and concave sides of dTMV assemblies.

## Discussion

Few artificial light harvesting systems have incorporated the two-dimensional fluidity afforded to natural light harvesting complexes by their embedment within a lipid membrane. Herein, we have adapted a well-characterized light harvesting model constructed from modified virus-like particles to facilitate their attachment to a planar lipid bilayer. This model was constructed from a newly reported dual-functional double-disk protein assembly consisting of 17 tandem dimer TMV subunits. By sequentially attaching a synthetic pigment and a His-tag-containing peptide to the dTMV surface, we have constructed a fully synthetic biomimetic model composed of discoidal protein arrays bound to chromophores and then associated with an SLB containing nickel-chelating lipids. TIRF microscopy measurements coupled with quantitative single molecule analysis enabled detailed characterization of the fluorescently labeled, bilayer-associated dTMV double disks and demonstrated their mobility on the SLB. The availability of the new dTMV construct represents a major advance in the use of viral capsids for templating pigments into light harvesting systems. Alongside the increased stability of its assembly state, dTMV also allows for asymmetric functionalization of opposing TMV complex faces and interior surfaces for the first time.

Whether all dTMV units are oriented with their N-terminal domains on one disk (e.g., the concave disk) and the C-terminal domain on the opposite disk (e.g., the flat disk) or whether they assemble at random, with the flat and concave disks each having a mixture of C- and N-terminal domains, remains an open question. The linker region between each domain was not well resolved in the crystal structure, and imaging of dTMV modified with biomolecules was similarly inconclusive. Because dTMV contains a flat and a concave disk, with the concave disk containing more adjacent monomer interface interactions than the flat disk, engineering at the intra-disk interfaces to remove or to add further residue interactions at only nTMV or cTMV′ may increase the likelihood of dTMV units being uniformly oriented within disk assemblies. Even if nTMV and cTMV′ are randomly oriented within double disk assemblies, there are residues that are more occluded in only the concave or only the flat disk as indicated by differences in their calculated solvent accessible surface areas (SASAs). As an example, residue Gln36 in the intracavity region between the disks of dTMV is involved in a hydrogen bonding network only present in the concave disk, and its mean SASA is 28.7±0.3 Å^2^ lower in the concave disk than in the flat disk (Table S1). Thus, a careful selection of protein modification site may target only a single side of a dTMV assembly even with a random orientation of dTMV units.

The protein modification strategies used herein have applications extending beyond the design of synthetic light harvesting models. The site-specific dual functionalization of proteins remains challenging but is extremely important for applications such as probing protein structure and function and developing precision therapeutics and diagnostic materials.(42) This work has demonstrated the site-selective conjugation of a dye and peptide at specific canonical amino acid residues engineered on a protein surface, adding to the toolbox of dual functionalization strategies. The covalent attachment of a His-tag to a protein surface site also has situational benefits over its conventional introduction as an N- or C-terminal tag during recombinant protein expression. Alongside protein purification, His-tags are widely used for protein attachment to surfaces.(43) However, genetically engineered His-tag fusion proteins can disrupt protein expression, folding, and enzymatic activity.(44–47) For proteins with buried N- and C-termini or those isolated from native organisms, expression with a His-tag may also be impractical. In contrast, introducing a single cysteine mutation at a convenient point on the protein surface as demonstrated herein or using a native surface-exposed cysteine can allow for post-expressional addition of a His-tag for downstream applications. This expands the library of proteins that can be associated with Ni-NTA-containing supported lipid bilayers and other surfaces.

The asymmetry of dTMV assemblies will allow for a more thorough interrogation of the role of the protein environment on chromophore excited state lifetimes and energy transfer in protein-based model light harvesting systems. Unlike *C*_*2*_-symmetric disk assemblies, the intracavity region of dTMV is narrower and can be selectively mutated on either the flat or concave surface to construct an asymmetric chromophore environment. In chromophore-labeled TMV complexes, these features of dTMV may allow for significant protein–chromophore coupling, a feature contributing to lengthened excited state lifetimes and high-efficiency energy transfer.(12) The solvent-exposed surface of dTMV is also easily accessible for functionalization, manifested by the successful step-wise conjugation of dyes and peptides described in this paper, leading to possibilities of coupling single dTMV assemblies to donor-acceptor chromophore pairs or other light harvesting proteins. The asymmetry of the complex lends itself to the directional attachment to a fluid lipid bilayer as shown in this work and could also be used to quantitate excited state energy transfer between mixed populations of dTMV disks that have been separately decorated with donor and acceptor chromophores and attached to an SLB. Such organization is reminiscent of the arrangement of LH1 and LH2 in photosynthetic membranes. The two-dimensional bilayer-attached dTMV model also provides a convenient platform for advanced spectroscopic and microscopic techniques such as Time-Resolved Ultrafast STED (TRUSTED)(24) and especially those requiring a supported surface such as single-particle fluorescence spectroscopy(20) to probe excited state energy migration along nanometer length scales.

## Materials and Methods

Protein purification, protein characterization methods, protein crystallization, X-ray diffraction data analysis, peptide synthesis, protein bioconjugation procedures, supported membrane preparation, TIRF microscopy, image analysis, and calculation of solvent-accessible surface area are described in details in SI Appendix.

## Acknowledgments

This work has been supported by the Director, Office of Science, Chemical Sciences, Geosciences, and Biosciences Division, of the U.S. Department of Energy under Contract No. DEAC02-05CH11231. X-ray diffraction data were collected at the Advanced Light Source (ALS) Beamline 8.2.1. The beamline is operated by the Berkeley Center for Structural Biology is supported in part by the Howard Hughes Medical Institute. The Advanced Light Source is a Department of Energy Office of Science User Facility under Contract No. DE-AC02-05CH11231. The ALS-ENABLE beamlines are supported in part by the National Institutes of Health, National Institute of General Medical Sciences, grant P30 GM124169. A. J. B. thanks a Chemical Biology Training Grant from the NIH (T32 GM066698) and the NSF Graduate Fellowship Program for financial support (DGE 1752814).

## References

1. T. Mirkovic, et al., Light Absorption and Energy Transfer in the Antenna Complexes of Photosynthetic Organisms. Chem. Rev. 117, 249–293 (2017).

2. J. N. Sturgis, J. D. Tucker, J. D. Olsen, C. N. Hunter, R. A. Niederman, Atomic Force Microscopy Studies of Native Photosynthetic Membranes †. Biochemistry 48, 3679–3698 (2009).

3. R. E. Blankenship, Molecular Mechanisms of Photosynthesis, R. E. Blankenship, Ed. (Blackwell Science Ltd, 2002) https://doi//.org/10.1002/9780470758472.

4. R. J. Cogdell, A. Gall, J. Köhler, The architecture and function of the light-harvesting apparatus of purple bacteria: from single molecules to in vivo membranes. Q. Rev. Biophys. 39, 227–324 (2006).

5. Y. Jiang, J. McNeill, Light-Harvesting and Amplified Energy Transfer in Conjugated Polymer Nanoparticles. Chem. Rev. 117, 838–859 (2017).

6. A. Adronov, J. M. J. Fréchet, Light-harvesting dendrimers. Chem. Commun., 1701–1710 (2000).

7. P. K. Dutta, et al., A DNA-directed light-harvesting/reaction center system. J. Am. Chem. Soc. 136, 16618–16625 (2014).

8. H. Park, et al., Enhanced energy transport in genetically engineered excitonic networks. Nat. Mater. 15, 211–216 (2016).

9. M. Delor, et al., Exploiting Chromophore–Protein Interactions through Linker Engineering To Tune Photoinduced Dynamics in a Biomimetic Light-Harvesting Platform. J. Am. Chem. Soc. 140, 6278–6287 (2018).

10. R. A. Miller, et al., Impact of Assembly State on the Defect Tolerance of TMV-Based Light Harvesting Arrays. J. Am. Chem. Soc. 132, 6068–6074 (2010).

11. K. B. Whaley, M. Sarovar, A. Ishizaki, Quantum entanglement phenomena in photosynthetic light harvesting complexes. 22nd Solvay Conf. Chem. 3, 152–164 (2011).

12. G. D. Scholes, G. R. Fleming, “Energy Transfer and Photosynthetic Light Harvesting” in (2005), pp. 57–129.

13. P. Akhtar, F. Görföl, G. Garab, P. H. Lambrev, Dependence of chlorophyll fluorescence quenching on the lipid-to-protein ratio in reconstituted light-harvesting complex II membranes containing lipid labels. Chem. Phys. 522, 242–248 (2019).

14. M. Yang, G. R. Fleming, Construction of kinetic domains in energy trapping processes and application to a photosynthetic light harvesting complex. J. Chem. Phys. 119, 5614–5622 (2003).

15. P. G. Adams, et al., Diblock Copolymer Micelles and Supported Films with Noncovalently Incorporated Chromophores: A Modular Platform for Efficient Energy Transfer. Nano Lett. 15, 2422–2428 (2015).

16. C. F. Calver, K. S. Schanze, G. Cosa, Biomimetic Light-Harvesting Antenna Based on the Self-Assembly of Conjugated Polyelectrolytes Embedded within Lipid Membranes. ACS Nano 10, 10598–10605 (2016).

17. R. A. Miller, A. D. Presley, M. B. Francis, Self-Assembling Light-Harvesting Systems from Synthetically Modified Tobacco Mosaic Virus Coat Proteins. J. Am. Chem. Soc. 129, 3104–3109 (2007).

18. Y.-Z. Ma, R. A. Miller, G. R. Fleming, M. B. Francis, Energy Transfer Dynamics in Light-Harvesting Assemblies Templated by the Tobacco Mosaic Virus Coat Protein. J. Phys. Chem. B 112, 6887–6892 (2008).

19. J. T. Groves, S. G. Boxer, Micropattern Formation in Supported Lipid Membranes. Acc. Chem. Res. 35, 149–157 (2002).

20. T. Kondo, W. J. Chen, G. S. Schlau-Cohen, Single-Molecule Fluorescence Spectroscopy of Photosynthetic Systems. Chem. Rev. 117, 860–898 (2017).

21. W. Y. C. Huang, H.-K. Chiang, J. T. Groves, Dynamic Scaling Analysis of Molecular Motion within the LAT:Grb2:SOS Protein Network on Membranes. Biophys. J. 113, 1807–1813 (2017).

22. W.-C. Lin, et al., H-Ras forms dimers on membrane surfaces via a protein–protein interface. Proc. Natl. Acad. Sci. 111, 2996–3001 (2014).

23. J. K. Chung, et al., K-Ras4B Remains Monomeric on Membranes over a Wide Range of Surface Densities and Lipid Compositions. Biophys. J. 114, 137–145 (2018).

24. S. B. Penwell, L. D. S. Ginsberg, R. Noriega, N. S. Ginsberg, Resolving ultrafast exciton migration in organic solids at the nanoscale. Nat. Mater. 16, 1136–1141 (2017).

25. J. T. Groves, N. Ulman, S. G. Boxer, Micropatterning Fluid Lipid Bilayers on Solid Supports. Science (80-.). 275, 651–653 (1997).

26. T. Lohmüller, et al., Single Molecule Tracking on Supported Membranes with Arrays of Optical Nanoantennas. Nano Lett. 12, 1717–1721 (2012).

27. L. Iversen, et al., Ras activation by SOS: Allosteric regulation by altered fluctuation dynamics. Science (80-.). 345, 50–54 (2014).

28. S. Yudovich, et al., Electrically controlling and optically observing the membrane potential of supported lipid bilayers. Biophys. J. 121, 2624–2637 (2022).

29. M. T. Dedeo, K. E. Duderstadt, J. M. Berger, M. B. Francis, Nanoscale Protein Assemblies from a Circular Permutant of the Tobacco Mosaic Virus. Nano Lett. 10, 181–186 (2009).

30. D. Liebschner, et al., Polder maps: Improving OMIT maps by excluding bulk solvent. Acta Crystallogr. Sect. D Struct. Biol. 73, 148–157 (2017).

31. B. Bhyravbhatla, S. J. Watowich, D. L. D. Caspar, Refined Atomic Model of the Four-Layer Aggregate of the Tobacco Mosaic Virus Coat Protein at 2.4-Å Resolution. Biophys. J. 74, 604–615 (1998).

32. X. Li, et al., Crystal Structure of a Four-Layer Aggregate of Engineered TMV CP Implies the Importance of Terminal Residues for Oligomer Assembly. PLoS One 8, 1–12 (2013).

33. M. T. Dedeo, K. E. Duderstadt, J. M. Berger, M. B. Francis, Nanoscale protein assemblies from a circular permutant of the tobacco mosaic virus. Nano Lett. (2010) https://doi//.org/10.1021/nl9032395.

34. M. T. Dedeo, K. E. Duderstadt, J. M. Berger, M. B. Francis, Nanoscale {Protein} {Assemblies} from a {Circular} {Permutant} of the {Tobacco} {Mosaic} {Virus}. Nano Lett. 10, 181–186 (2010).

35. M. A. Bruckman, et al., Role of Hexahistidine in Directed Nanoassemblies of Tobacco Mosaic Virus Coat Protein. ACS Nano 5, 1606–1616 (2011).

36. C. S. Mogilevsky, et al., Synthesis of Multi-Protein Complexes through Charge-Directed Sequential Activation of Tyrosine Residues. J. Am. Chem. Soc. 143, 13538–13547 (2021).

37. A. M. Marmelstein, et al., Tyrosinase-Mediated Oxidative Coupling of Tyrosine Tags on Peptides and Proteins. J. Am. Chem. Soc. 142, 5078–5086 (2020).

38. J. C. Maza, et al., Enzymatic Modification of N-Terminal Proline Residues Using Phenol Derivatives. J. Am. Chem. Soc. 141, 3885–3892 (2019).

39. J. A. Nye, J. T. Groves, Kinetic Control of Histidine-Tagged Protein Surface Density on Supported Lipid Bilayers. Langmuir 24, 4145–4149 (2008).

40. Y. Kaizuka, J. T. Groves, Structure and Dynamics of Supported Intermembrane Junctions. Biophys. J. 86, 905–912 (2004).

41. J. C. Maza, et al., Tyrosinase-Mediated Synthesis of Nanobody–Cell Conjugates. ACS Cent. Sci. 0 (2022).

42. L. Xu, S. L. Kuan, T. Weil, Contemporary Approaches for Site-Selective Dual Functionalization of Proteins. Angew. Chemie Int. Ed. 60, 13757–13777 (2021).

43. R. Wieneke, R. Tampé, Multivalent Chelators for In Vivo Protein Labeling. Angew. Chemie Int. Ed. 58, 8278–8290 (2019).

44. S. Zhu, et al., A simple and effective strategy for solving the problem of inclusion bodies in recombinant protein technology: His-tag deletions enhance soluble expression. Appl. Microbiol. Biotechnol. 97, 837–845 (2013).

45. D. Zhao, Z. Huang, Effect of His-Tag on Expression, Purification, and Structure of Zinc Finger Protein, ZNF191(243-368). Bioinorg. Chem. Appl. 2016, 8206854 (2016).

46. L. Meng, et al., Effects of His-tag on Catalytic Activity and Enantioselectivity of Recombinant Transaminases. Appl. Biochem. Biotechnol. 190, 880–895 (2020).

47. E. A. Woestenenk, M. Hammarström, S. van den Berg, T. Härd, H. Berglund, His tag effect on solubility of human proteins produced in Escherichia coli: a comparison between four expression vectors. J. Struct. Funct. Genomics 5, 217–229 (2004).

