## Supporting Information for "A membrane-associated light harvesting model is enabled by functionalized assemblies of gene-doubled TMV proteins"

**This PDF file includes:**

Supporting text
Figures S1 to S12
Table S1
Legends for Movies S1 to S4
SI References

**Other supporting materials for this manuscript include the following:**

Movies S1 to S4

### Supporting Information Text

**General methods and materials.** Unless otherwise noted, all chemicals and solvents were of analytical grade and were received from commercial sources. The phospholipids 1,2-dioleoyl-sn-glycero-3-phosphocholine (DOPC), 1,2-dioleoyl-sn-glycero-3-[(*N*-(5-amino-1-carboxypentyl)iminodiacetic acid)succinyl] nickel salt (Ni-NTA-DOGS), and 1,2-dioleoyl-sn-glycero-3 phosphoethanolamine-*N*-[4-(*p*-maleimidomethyl)cyclohexane-carboxamide] (MCC-DOPE) sodium salt for preparation of supported membranes were purchased from Avanti Polar Lipids (Alabaster, AL) as chloroform solutions. Alexa Fluor 647 NHS ester was purchased from Sigma-Aldrich (St. Louis, MO). Water (dd-$H_2O$) used as reaction solvent was deionized using a Barnstead NANOpure purification system (Thermo Fisher, Waltham, MA). All oligonucleotides were purchased from Integrated DNA Technologies (Coralville, IA). Protected amino acids and resins for solid phase peptide synthesis were obtained from Novabiochem (Merck KGaA, Darmstadt, Germany). Spin concentration was performed using 100,000 or 30,000 molecular weight cutoff spin concentrators from Millipore (Burlington, MA). Dialysis was performed with either Slide-A-Lyzer Dialysis Cassettes (Pierce, Rockford, Il) for small volume samples or dialysis tubing (Fisherbrand, Pittsburgh, PA) for large volume samples.

**Mass spectrometry.** Protein and small molecules were analyzed using liquid chromatography (1200 series, Agilent Technologies, USA) that was connected in line with an Agilent 6224 Time-of-Flight (TOF) mass spectral system equipped with a Turbospray ion source. Protein samples were run with a Proswift RP-4H column (Dionex, USA). Protein mass reconstruction was performed on the charge ladder with Mass Hunter software (Agilent Technologies, USA).

**Steady-state Spectroscopy.** UV-Vis absorption measurements were conducted on a Cary UV-Vis 100 spectrophotometer (Agilent, USA). Steady-state fluorescence measurements were obtained on a Fluoromax-4 spectrofluorometer (Horiba Scientific, USA). Slit widths were set to 1.0 nm for both excitation and emission. Protein concentration was routinely determined by UV/Vis analysis on a Nanodrop 1000 instrument (Nanodrop, USA) by monitoring absorbance at 280 nm.

**High Performance Liquid Chromatography.** HPLC was performed on Agilent 1260 Infinity Series HPLC Systems (Agilent, USA). Sample analysis for all HPLC experiments was achieved with an in-line diode array detector (DAD) and an in-line fluorescence detector (FLD). Size exclusion chromatography (SEC) was performed using a Polysep-GFC-P-5000 column (4.6 x 250 mm) (Phenomenex, USA) at 1.0 mL/min using a mobile phase of 10 mM sodium phosphate buffer, pH 7.2.

**Gel Analysis.** Sodium dodecyl sulfate-polyacrylamide gel electrophoresis (SDS-PAGE) was carried out in a Mini cell tank apparatus (Life Technologies, Carlsbad, CA), using NuPAGE™ Novex™ 4-12% Bis-Tris Protein Gels (Life Technologies). The sample and electrode buffers were prepared according to the suggestions of the manufacturer. All protein electrophoresis samples were heated for 5-10 min at 95 °C in the presence of 1,4-dithiothreitol (DTT) to ensure the reduction of disulfide bonds. Gels were run for 30 min at 200 V to separate the bands. Commercially available markers (Bio-Rad) were applied to at least one lane of each gel for the assignment of apparent molecular masses. Visualization of protein bands was accomplished by staining with Coomassie Brilliant Blue R-250 (Bio-Rad, Hercules, CA). Gel imaging was performed on a Gel Doc (Bio-Rad, Hercules, CA).

**Transmission Electron Microscopy.** TEM analysis of dTMV and its conjugates with peptide and nanobody was carried out at the Berkeley Electron Microscope Lab with an FEI Tecnai 12 transmission electron microscope with 100 kV accelerating voltage. Samples were prepared for analysis by applying 5 µL analyte solution (10 µM protein in 10 mM NaPhos pH 7.2) to carbon-coated copper grids for 2 min. The sample application was followed by rinsing in 4 x 10 µL of 1% uranyl acetate solution, leaving the sample in the last droplet for 1 min before wicking away the excess solution.

**Generation of dTMV mutants***.* The gene for the permuted TMV coat protein (dTMV) was cloned into an *E. coli* expression vector (pBAD, Life Technologies). The starting plasmid contains two mutations S123C and S123′C. The QuickChange II Site-Directed Mutagenesis Kit (Agilent Technologies, USA) was used to generate introduce mutations C123S, C123′S, C123K, C123′K and N-terminus insertion using the following sets of primers:
C123S:
Sense: 5' GCCACCGTGGCGATTCGCAGTGCCATCAATAACC 3'
Antisense: 5' GGTTATTGATGGCACTGCGAATCGCCACGGTGGC 3'
C123′S:
Sense: 5' CGCAACGGTGGCCATAAGGAGCGCGATAAATAATTTAATAGTAG 3'
Antisense: 5' CTACTATTAAATTATTTATCGCGCTCCTTATGGCCACCGTTGCG 3'
C123K:
Sense: 5' CCACCGTGGCGATTCGCAAGGCCATCAATAACCTGATTGTGG 3'
Antisense: 5' CCACAATCAGGTTATTGATGGCCTTGCGAATCGCCACGGTGG 3'
C123′K:
Sense:
5' GGTGGCCATAAGGAAGGCGATAAATAATTTAATAGTAGAATTGATCAGAGG 3'
Antisense:
5' CCTCTGATCAATTCTACTATTAAATTATTTATCGCCTTCCTTATGGCCACC 3'
Insertion of A at N-terminus:
Sense: 5' GGAGGAATTAACCATGGCCGGCAGCTATAGCATTACCACCCC 3'
Antisense: 5' GGTAATGCTATAGCTGCCGGCCATGGTTAATTCCTCCTGTTAGC 3'

**Protein expression and purification.** DH10B competent cells were transformed with the plasmids containing the dTMV variants described above. Colonies were selected for inoculation in Terrific Broth with 100 μg/L ampicillin at 37 °C. When cultures reached an optical density of 0.6 to 0.8, 0.01% arabinose was added. After growing for 20 h at 20 °C, the cells were harvested by centrifugation (8000 rpm, 15 min) and the cell pellet was stored at -20 °C. Cells were resuspended in 20 mL lysis buffer (20 mM triethanolamine [TEA] pH 7.2) supplemented with benzonase and 2 mM $MgCl_2$. Cells were lysed by sonication with a 2 s on, 4 s off cycle for a total of 10 min using a standard disruptor horn at 60% amplitude (Branson Ultrasonics, Danbury, CT). The resulting lysate was cleared at 14,000 rpm for 30 min. A saturated solution of ammonium sulfate was added to the supernatant to reach the final concentration of 15%. The mixture was rotated for 10 min at 4 °C to allow the complete protein precipitation. The precipitated protein was then collected at 10,000 rpm for 30 min and then resuspended in low salt buffer (20 mM TEA, pH 7.2). The resulting protein solution was dialyzed against the buffer to remove the residual ammonium sulfate before loading onto a DEAE column and purified with a 0–300 mM NaCl gradient elution in buffer (20 mM TEA, pH 7.2). Purity was confirmed by SDS-PAGE and ESI-TOF MS. Pure fractions were pooled, and fractions containing desired TMV in addition to impurities were further purified using HiPrep™ 26/60 Sephacryl® S-500 HR column (GE Healthcare, USA).

**Protein crystallization.** Purified dTMV was buffer exchanged into HEPES buffer (25 mM HEPES, 50 mM NaCl, 2 mM DTT, pH 7.2) and concentrated to 10–13 mg/mL. Crystals were formed by mixing 0.2 μL of the protein with 0.2–0.3 μL crystallization buffer (100 mM Bis-Tris, pH 6.5, 45 % (v/v) polypropylene glycol P400) in 96 well sitting-drop plates (Intelli-plate 96-3 LVR) before incubation over 1-3 weeks at 20±0.5 °C. The crystals were harvested in mother liquor supplemented with 5% (v/v) glycerol before being flash frozen in liquid nitrogen.

**X-ray diffraction data collection and model refinement.** X-ray diffraction data were collected at the Advanced Light Source (ALS) Beamline 8.2.1 under cryogenic conditions. X-ray diffraction data were indexed and scaled with XDS (1) before merging in AIMLESS (2) with an applied resolution cut-off of 2.8 Å based on the half-dataset correlation coefficient $CC^{1/2}$ (3). Phasing was done using molecular replacement with the reported wild type TMV monomer structure (PDB code: 1EI7). The model was iteratively built in COOT (4) and refined using PHENIX.refine (5, 6) with rigid body, XYZ (reciprocal-space), group B-factors, secondary structure, non-crystallographic symmetry, stereochemistry, and grouped atomic displacement parameters (ADP) restraints.

**General procedure for peptide synthesis.** Solid-phase peptide synthesis was performed following an established literature procedure with minimal modifications (7). Side chain protecting groups used were: His (Trt), and Ser (*t*Bu). The resin linker used was benzyloxybenzyl alcohol (Wang) polystyrene. Synthesis was accomplished manually, using 5 equiv. of amino acids in dimethylformamide (DMF) with *O*-(1*H*-6-Chlorobenzotriazole-1-yl)-1,1,3,3-tetramethyluronium hexafluorophosphate (HCTU, Novabiochem, Merck KGaA, Darmstadt, Germany) as the coupling reagent with 10 equiv. *N*,*N*-diisopropylethylamine (DIPEA, Sigma-Aldrich, St. Louis, MO) as an additive. Once all amino acids had been coupled to the peptide on the resin, the N-terminal Fmoc group was removed using 20% piperidine (Sigma-Aldrich, St. Louis, MO) in DMF. Peptides were cleaved from resin using a cocktail of 95% trifluoroacetic acid (TFA, EMD Millipore, Billerica, MA), 2.5% water, and 2.5% triisopropylsilane (TIPS, Sigma-Aldrich, St. Louis, MO). Crude peptides were precipitated in cold diethyl ether, analyzed, and further purified if needed by reversed-phase HPLC, and lyophilized before use.

| Peptide Data Table | | | |
|---|---|---|---|
| **Name** | **Sequence** | **MW (calculated)** | **MW (found)** |
| $H_6Y$ | HHHHHHY | 1004.0 | 1003.4 |
| $H_6SG_4Y$ | HHHHHHSGGGGY | 1319.3 | 1318.5 |

**General procedure for labeling of dTMV with AF647 NHS ester.** The proteins were first exchanged into the reaction buffer (100 mM sodium phosphate, pH 8.5). To 100 µL of dTMV (100 µM) was added 2 equiv. of AF647. The reaction mixture was briefly agitated and then incubated in 1.5 mL Eppendorf tubes at room temperature with an aluminum foil cover. After 15 min, the reactions were quenched by 1 mM hydroxylamine. The crude reaction was purified with a NAP-5 Sephadex G-25 column (GE Healthcare, USA) followed by purification using a Polysep-GFC-P-5000 column (4.6 x 250 mm) (Phenomenex, USA) to remove the unreacted chromophores. The fractions that showed absorption at 280 nm were combined and subjected to spin concentration using 100 kDa MWCO filters. The protein conjugates were analyzed with LC-MS and HPLC-SEC for assessment of conjugation level, purity, and validation of assembly state.

**General procedure for labeling of dTMV with $H_6Y$ or $H_6SG_4Y$ peptide.** The proteins were first exchanged into the reaction buffer (10 mM sodium phosphate, pH 7.2). To 100 µL of peptide (500 µM) was added 400 nM tyrosinase from *Agaricus bisporus* (abTyr), followed by 25 µM protein. The reaction mixture was briefly agitated and then incubated in 1.5 mL Eppendorf tubes at room temperature with an aluminum foil cover. After 2 h, excess peptide and abTyr were removed via repeated centrifugation through 100 kDa molecular weight cutoff filters and used directly for TEM imaging or further purified for fluorescence imaging using a Polysep-GFC-P-5000 column (4.6 x 250 mm). The fractions in the size range containing TMV disk stacks were combined and subjected to spin concentration using 100 kDa MWCO filters. The protein conjugates were analyzed with MS and HPLC-SEC for assessment of conjugation level, purity, and validation of assembly state.

**Procedure for labeling of Bovine Serum Albumin (BSA) with AlexaFluor 647 (AF647) NHS ester.** BSA lyophilized powder (Sigma-Aldrich, St. Louis, MO) was dissolved just before use into the reaction buffer (100 mM sodium phosphate, pH 8.5) at a concentration of 100 µM. To 50 µL of BSA (100 µM) was added 2 equiv. of AF647. The reaction mixture was briefly agitated and then incubated in 1.5 mL Eppendorf tubes at room temperature with an aluminum foil cover. After 1 h, the reactions were quenched by 1 mM hydroxylamine. The crude reaction was purified with a NAP-5 Sephadex G-25 column followed by purification using a Polysep-GFC-P-5000 column (4.6 x 250 mm) to remove the unreacted chromophores. The fractions that showed absorption at 280 nm were combined and subjected to spin concentration using 30 kDa MWCO filters. The protein conjugates were analyzed with LC-MS and HPLC-SEC for assessment of conjugation level and purity.

**Procedure for labeling the BSA-AF647 conjugate with $H_6$Y peptide.** BSA-AF647 was first exchanged into the reaction buffer (10 mM sodium phosphate, pH 7.2). To 100 µL of $H_6$Y peptide (200 µM) was added 400 nM tyrosinase from *Agaricus bisporus* (abTyr), followed by 100 µM protein. The reaction mixture was briefly agitated and then incubated in 1.5 mL Eppendorf tubes at room temperature with an aluminum foil cover. After 2.5 h, excess peptide and abTyr were removed via repeated centrifugation through 30 kDa molecular weight cutoff filters followed by purification using a Polysep-GFC-P-5000 column (4.6 x 250 mm). The fractions that showed absorption at 280 nm were combined and subjected to spin concentration using 30 kDa MWCO filters. The protein conjugates were analyzed with MS and HPLC-SEC for assessment of conjugation level and purity.

**Procedure for constructing dTMV-nbHER2$_{Tyr}$ conjugate.** The proteins were first exchanged into the reaction buffer (10 mM sodium phosphate, pH 7.2). To 100 µL of nbHER2$_{Tyr}$ (50 µM) was added 400 nM tyrosinase from *Agaricus bisporus* (abTyr), followed by 25 µM dTMV-S123C–S123′K. The reaction mixture was briefly agitated and then incubated in 1.5 mL Eppendorf tubes at room temperature with an aluminum foil cover. After 2 h, excess nbHER2$_{Tyr}$ and abTyr were removed via repeated centrifugation through 100 kDa molecular weight cutoff filters. The protein conjugates were analyzed with MS and HPLC-SEC for assessment of conjugation level, purity, and validation of assembly state.

**Supported membrane preparation.** Supported membranes were formed on glass coverslips as described elsewhere (8). Briefly, 25 mm #1.5 thickness round glass coverslips (Thomas Scientific, Swedesboro, NJ) were ultrasonicated for 30 min in 50:50 isopropyl alcohol/water, rinsed thoroughly in Milli-Q water (MilliporeSigma, Billerica, MA), etched for 5 min in freshly prepared piranha solution (3:1 sulfuric acid/hydrogen peroxide), and again rinsed thoroughly in Milli-Q water. Coverslips were then assembled in Attofluor chambers (Invitrogen, Waltham, MA) that had been cleaned in 50:50 isopropyl alcohol/water, rinsed thoroughly in Milli-Q water, and dried. Bilayers were formed by rupturing small unilamellar vesicles (SUVs) on the cleaned glass substrate. SUVs for bilayer formation were prepared by mixing a molar ratio of 98% DOPC and 2% Ni-NTA-DOGS in chloroform in a round bottom flask, drying the lipid mixture with a rotovap, and resuspending the mixture to 0.5 mg/mL in Milli-Q water.

For bilayers including maleimide-headgroup lipids, the lipid mixture contained molar ratios of 95% DOPC, 2% Ni-NTA-DOGS, and 3% MCC-DOPE. The suspension of lipids in water was then sonicated with a probe sonicator (Analis Scientific Instruments, Namur, Belgium) for a total of 1 min using pulses of 15 s at 32% amplitude with 10 s pauses while sitting in an ice bath to prevent the lipid suspension from warming. Sonicated solutions were then centrifuged at 21,000 g for 20 min at 4 °C to remove titanium particles from the sonicator and lipid aggregates. SUV solutions in water were mixed 1:1 with PBS and 300 µL of this solution was added to each assembled Attofluor chamber and incubated for 35 min. The membrane was washed with PBS and then incubated in imaging buffer (20 mM HEPES, 137 mM NaCl, 1 mM $CaCl_2$, 2 mM $MgCl_2$, 5 mM KCl, 0.7 mM $Na_2HPO_4$, 6 mM D-glucose, and 0.2% w/v bovine serum albumin (BSA)) for 30 min to block bilayer defects and prevent nonspecific interactions of fluorescent proteins of interest with the surface. dTMV was diluted in an imaging buffer to 4x the desired working concentration, then diluted 1:4 into the Attofluor chambers and incubated for 30–35 min. The solution incubation concentrations of dTMV proteins were 15-40 nM for experiments requiring high protein density on the bilayer and and 100-500 pM for single particle tracking experiments.

**TIRF microscopy.** TIRF experiments were performed on a motorized inverted microscope (Nikon Eclipse Ti-E; Technical Instruments, Burlingame, CA) equipped with a motorized Epi/TIRF illuminator (Nikon), Lumen Dynamics X-Cite® 120LED Fluorescence Illumination System (Excelitas Technologies, Waltham, MA, USA), Perfect Focus System (Nikon), and a motorized stage (Applied Scientific Instrumentation MS-2000, Eugene, OR). A laser launch with 488, 561, and 640 nm (Coherent OBIS, Santa Clara, CA) diode lasers was controlled by an OBIS Scientific Remote (Coherent) and aligned into a fiber launch custom built by Solamere Technology Group (Salt Lake City, UT). A dichroic beamsplitter (z488/647rpc; Chroma Technology, Bellows Falls, VT) reflected the laser light through the objective lens, and fluorescence images were recorded using an electron multiplying (EM)-CCD (iXon 897DU; Andor, South Windsor, CT) after passing through a laserblocking filter (Z488/647M; Chroma Technology). Exposure times, multidimensional acquisitions, and time-lapse periods for all experiments were set using Micro-Manager (9). A transistor-transistor logic signal from the appropriate laser triggered the camera exposure. The laser intensities were measured at the sample for each experiment day so that a constant laser intensity is used for each type of imaging across different days.

dTMV density on bilayers was imaged using 20 ms exposure time, 0.7-1.2 mW power at the sample, and a camera setting of 500 gain. Imaging of AF647-labeled dTMV diffusion on the supported membrane was performed with a streaming acquisition of 20 ms exposure time at 5.2 mW power at the sample and 1000 gain for 50 frames. Oxygen scavengers were not used to minimize photobleaching in order to analyze the number of discrete step photobleaching events along particle trajectories. Images for the dTMV single particle intensity distribution were taken with the same acquisition settings as for the diffusion data. For FRAP measurements, TIRF images before and after bleaching were collected every 20 s at 0.2 mW power and 20 ms exposure time. The bleaching was performed in TIRF, closing the front aperture to define the bleached spot, with 11 mW power continuously for 1.5 s. All imaging was done at 24 °C.

**Image Analysis.** Image analysis was performed in MATLAB (MathWorks, Natick MA) using custom scripts, available upon request.

*Quantitation of particle density on SLBs.* An intensity vs. density calibration curve was built from at least 20 micrographs each of 4 bilayers at multiple dTMV densities for which particles were countable, using the same imaging conditions (20 ms, 1.2 mW at sample, 500 gain) for each sample. Particles in a 28 μm x 28 μm area of even illumination were counted using TrackMate, with the particle diameter and intensity threshold set by eye then applied uniformly across the data sets. The mean intensity of the same area was measured for each bilayer. The slope and y-intercept of this linear intensity vs. density calibration curve was used to calculate the density of dTMV on high-density bilayers. Images of high-density bilayers were taken with decreased illumination intensity, but as fluorescence intensity scales linearly with illumination intensity, this change in imaging conditions was easily accounted for to calculate bilayer density.

*FRAP analysis.* FRAP data were normalized according to the workflow described in Carnell, M et al. (10). Briefly, a circular FRAP region fully contained in the bleached area was defined, and at each timepoint *t*, the mean intensity in this area, FRAP(t), was subtracted by intensity in that area immediately after bleaching ($FRAP_{bleach}$). This quantity was normalized to the pre-bleach intensity of the FRAP area ($FRAP_{pre\text{-}bleach}$) and the intensity of a reference area at time *t* ($Ref_{pre\text{-}bleach}$/ref(t)). The fully normalized FRAP recovery trace was calculated as,

$$Full\,norm(t) = \frac{Ref_{pre-bleach}}{ref(t)} \cdot \frac{FRAP(t) - FRAP_{bleach}}{FRAP_{pre-bleach} - FRAP_{bleach}}$$

The resulting recovery trace was fit to a diffusion-limited recovery model,

$$Full\,norm(t) = A \cdot e^{\frac{2\tau_D}{t}}\left(I_0\left(\frac{2\tau_D}{t}\right) + I_1\left(\frac{2\tau_D}{t}\right)\right)$$

where

$$\tau_D = \frac{w^2}{4D}$$

and where A is the plateau intensity, w is the bleach area radius, D is the diffusion coefficient, and $I_0()$ and $I_1()$ are modified Bessel functions of the first kind. This model assumes bleaching occurs instantaneously, bleaching has a step-function profile, recovery is diffusion limited, diffusion is lateral and equal in all directions, and there is a single diffusing population with a single diffusion coefficient. Our experiment generally meets all assumptions except for the last, but a fit can allow us to extract a mean diffusion coefficient from the highly heterogeneous sample.

*Single particle diffusion analysis.* Particles in a 28 µm x 28 µm area of even illumination were tracked using TrackMate using the Difference of Gaussians spot detector, a particle diameter of 0.4 µm, threshold of 300, and using the median filter and sub-pixel localization. The simple LAP tracker was used to link particle trajectories, using a maximum linking distance of 0.4 µm and not allowing gaps in tracks. The data from TrackMate was then exported to MATLAB for analysis and visualization. The diffusion coefficient for individual tracks was calculated from the mean square displacement between adjacent frames along the track. The step size distribution of 2,281 particles was compiled from particle localizations from every third frame along particle trajectories longer than 10 frames. This 60 ms time interval minimizes the effect of particle localization errors, which can introduce artifacts with shorter time intervals (11). Due to the highly heterogeneous diffusion within the sample, we chose not to fit the step size distribution to a model with a specific number of diffusing species.

*Quantitation of single particle intensities.* TMV particles presented on bilayers at low density (0.05 $\mu m^{-2}$) were localized in 28 µm x 28 µm areas of even illumination using TrackMate using parameters stated above. Total particle intensity was determined by integrating a 7 pixel x 7 pixel (0.75 µm x 0.75 µm) area centered around the particle and subtracting the intensity of an average background area of the same size. These intensities were then calibrated to the intensity of a single fluorophore, determined by integrating a 7 pixel x 7 pixel area around single emitters (12), to obtain a distribution of the number of fluorophores bound to TMV particles.

*Step photobleaching analysis.* Tracks obtained for single particle diffusion analysis were analyzed for the number of step photobleaching events. Only tracks of particles that were reliably tracked throughout the acquisition and fully bleach within the acquisition were analyzed to ensure that the number of fluorophores attached to each disk was measured accurately. The particle intensity in each frame was determined as stated above. Upon particle bleaching, an additional 10 frames (or until the end of the acquisition) were analyzed for total background-subtracted intensity at the last identified particle location and appended to the particle intensity trace. The intensity trace was analyzed using a Bayesian change point algorithm (13) to determine the number of change points, or fluorophore bleaching events, in the particle trajectory. All intensity traces and identified change points were verified manually.

**Procedure for generating cpTMV homology model.** For visualization purposes, a homology model of cpTMV containing all amino acid residues was generated based on the crystal structure of cpTMV (PDB code 3KML) (14) and the crystal structure of wtTMV (PDB code 1EI7) (15), both obtained from the Protein Data Bank. Initial structural preparation was conducted using The Pymol Molecular Graphics System, Version 2.4.2. First, a monomer of wtTMV was superimposed on a monomer of cpTMV, and the residues resolved in the crystal structure of wtTMV but not cpTMV (residues 92–110 of wtTMV and 1–12, 154–161 of cpTMV) were fused to the unresolved N- and C-termini of cpTMV. The bond between residues 99 and 100 of wtTMV was then cleaved *in silico* and an N-terminal glycine was added to produce the N- and C-termini of cpTMV. Following this, cpTMV was symmetry expanded to create the double disk assembled structure consisting of two $C_2$-symmetric disks, each containing 17 monomers. The Schrödinger Maestro package (version 2022-1) (16) was used for subsequent structural preparation and molecular dynamics simulations. The Desmond system builder was used to solvate the double disk structure in an orthorhombic box with periodic boundaries at 10 Å from the protein of water molecules described using the TIP3P model,(17, 18) neutralized with sodium ions, with the addition of 150 mM NaCl in an OPLS4 force field.(19) A molecular dynamics simulation was performed with an NPT ensemble of T = 300 K, P = 1 bar, a Coulombic cutoff radius of 9.0 Å, and a 100 ps simulation time with sampling time of 5 ps. This short simulation time was chosen to relax sidechain and solvent interactions for the protein representations shown herein, but not alter the overall quaternary structure of the assembly. The pressure control was applied using the Martyna-Tobias-Klein barostat method(20) with a 2.0 ps relaxation time, and the temperature control was applied using the Nosé-Hoover thermostat method with a 1.0 ps relaxation time.(21) The trajectory was analyzed, and the lowest energy frame was used as the homology model.

**Procedure for obtaining solvent-accessible surface area (SASA) of dTMV residues.** The SASA of each amino acid residue of dTMV was obtained using the crystal structure of dTMV reported herein as an input structure. The Schrödinger Bioluminate package (22) was used to perform a Residue Analysis of all resolved residues present on all 17 tandem dimers of the dTMV crystal structure to obtain the SASA of each residue. The corresponding residues of each tandem dimer were averaged, and the averaged value of each residue of nTMV and the corresponding cTMV′ residue were compared. The resulting values are shown in Table S1. Statistical analyses were performed in Microsoft Excel for Mac, Version 16.62.

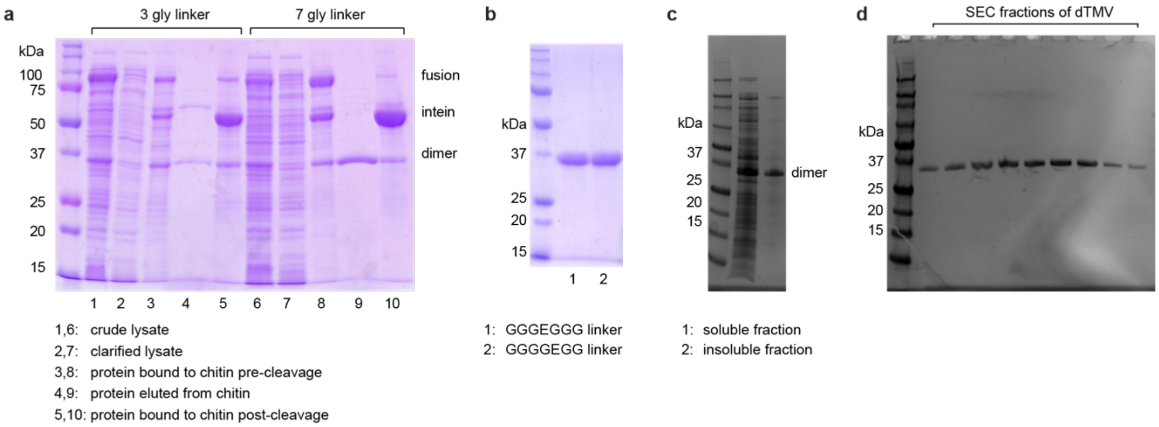


**Fig. S1.** Testing linker length and expression conditions of dTMV. (a) Test expressions of a TMV tandem dimer fused with intein from a pTYB1 vector show that the best expression conditions were using a 7-glycine linker between monomer subunits. (b) The addition of a glutamate residue to the amino acid linker between monomer subunits did not affect protein expression with the glutamate in the fourth (lane 1) or fifth (lane 2) position of the linker. (c) Expression of dTMV with the GGGEGGG linker from the pBAD vector, rather than as an intein fusion, also resulted in an acceptable level of expression. (d) Purification from the soluble fraction after expression from the pBAD vector was achieved via precipitation of dTMV with ammonium sulfate, followed by resolubilization and ion exchange chromatography followed by size exclusion chromatography (SEC). The pure fractions collected following SEC purification are shown.

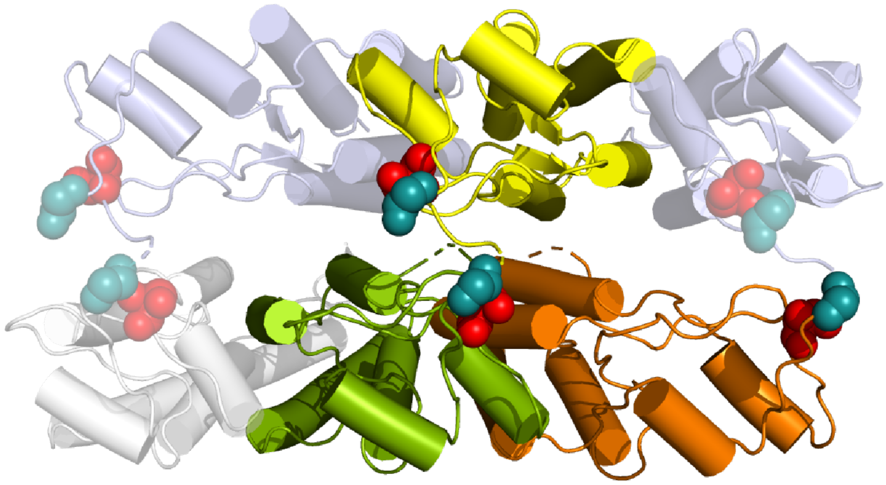

**Fig. S2.** Determination of the location of nTMV and cTMV′ of dTMV within disk assemblies. Six adjacent monomers within the double disk assembly are shown for clarity. The first residue resolved in the N-terminal region, Ser1, is shown as red spheres, and the last residue resolved in the C-terminal region, Gly155′, is shown as cyan spheres. There are 10 unresolved residues, including the 7-amino-acid GGGEGGG linker as well as 3 residues at the C-terminus of nTMV. The estimated length of a linearized 10-amino-acid peptide is roughly 35 Å. The linear distance between the N- and C-termini of adjacent monomers within a single disk in the dTMV assembly, shown in green and orange above, is 33.5 Å, which is close to the maximum length limit of the linkage residues. Modeling in 10-amino-acid linkers to fuse green and orange monomers without causing clashes with existing protein structures did not yield any reasonable results. This indicates that nTMV and cTMV′ of a single dTMV unit cannot occupy locations adjacent to one another on the same disk. Instead, nTMV and cTMV′ of a single dTMV unit must always occupy positions in opposing disks, as demonstrated by the yellow and green colored subunits.

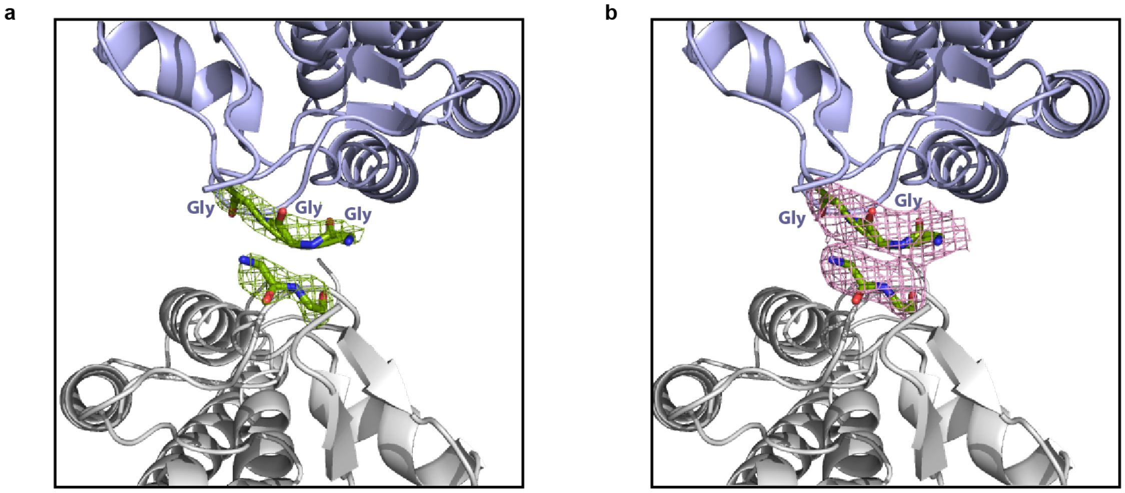


**Fig. S3.** Electron density maps around the hypothesized linker position. (a) The positive $mF_O–DF_C$ map (green meshed line) and (b) the polder map (pink meshed line) for the potential linker position are contoured at the 3.0 σ level. Three amino acids of the linker (GGG, colored green) are modeled extending from the N-terminus of the flat monomer (colored purple). Two amino acids of the linker (GG, colored green) are modeled extending from the N-terminus of the concave monomer (colored grey).

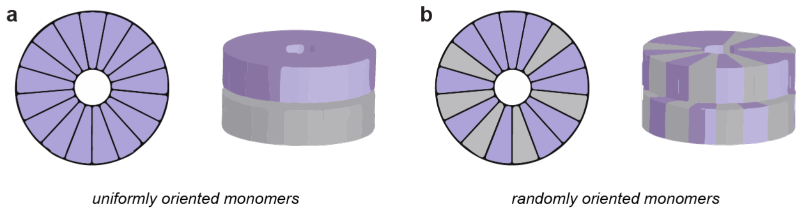


**Fig. S4.** Potential orientation of tandem dimers within dTMV assemblies. (a) A schematic shows one possibility of tandem dimers arranged in a uniform orientation within double disk assemblies, with the purple section being the N-terminal portion of the tandem dimer (nTMV) and the gray section being the C-terminal portion (cTMV′). (b) A schematic shows the second possibility of tandem dimers arranged in a random orientation within double disk assemblies, with the purple section corresponding to nTMV and the gray portion corresponding to cTMV′.

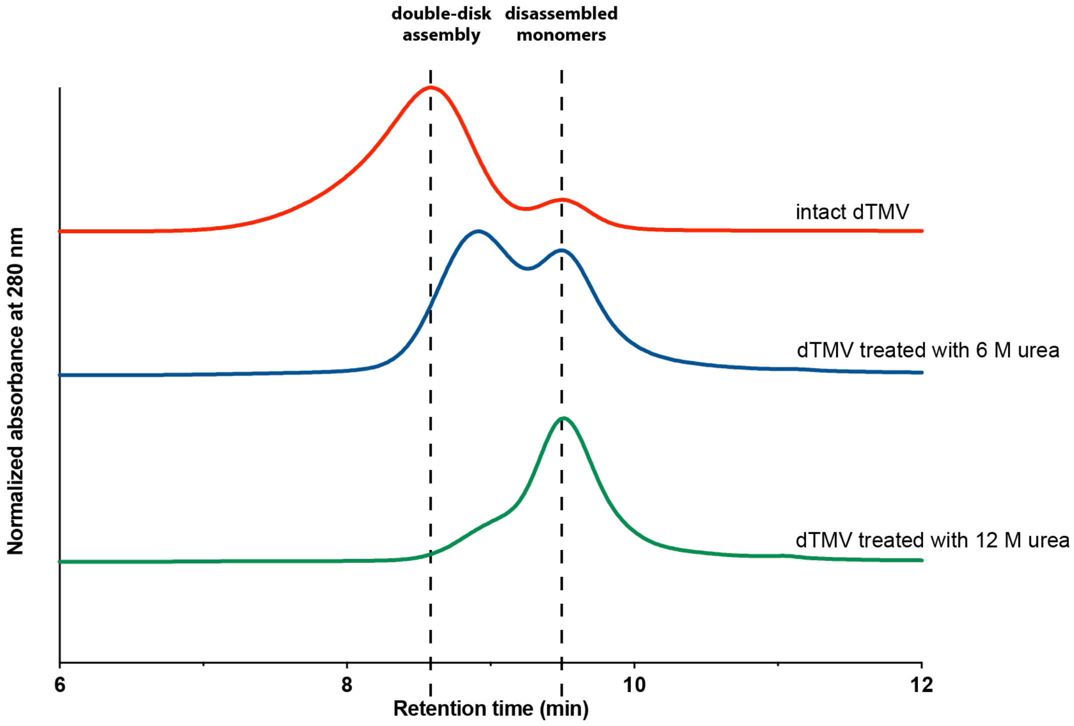


**Fig. S5.** Disassembly of dTMV characterized by size exclusion chromatography (SEC). A shift in retention time compared to the SEC profile of the intact dTMV (red trace) indicated the assembly state change. A complete disassembly took place when treated with 12 M urea (green trace). Partial disassembly resulting in a mixture of different assembly species were observed at a lower concentration of urea (blue trace).

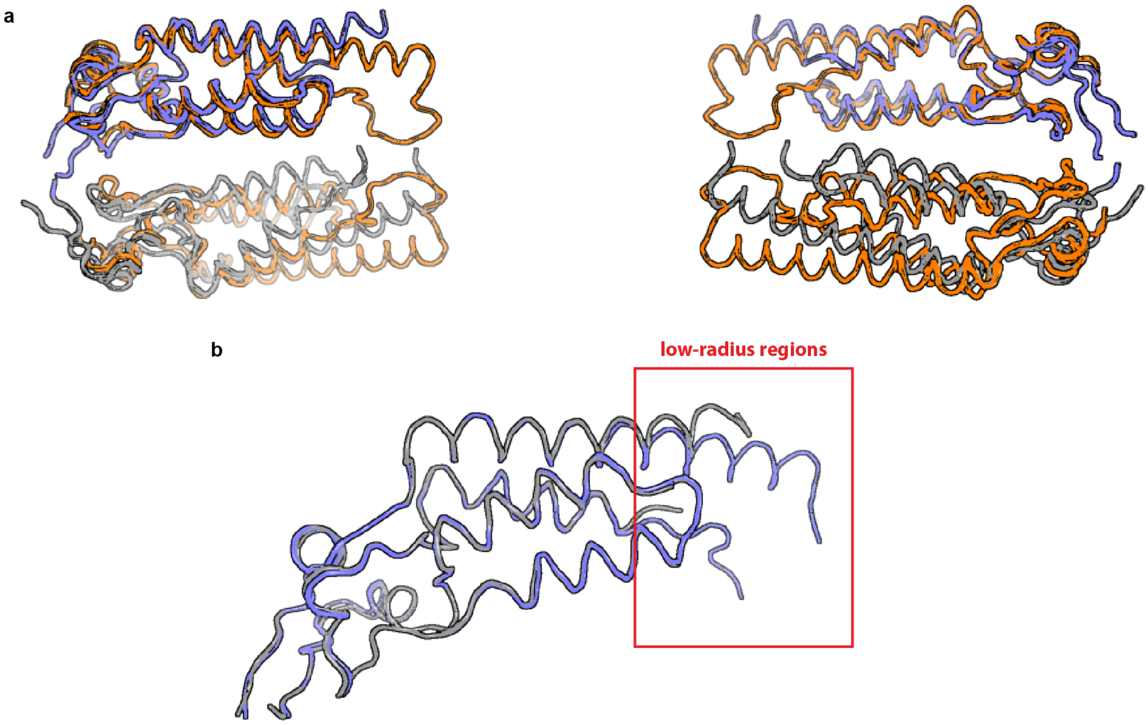


**Fig. S6.** Superimposition of TMV backbones. (a) A structural comparison between the dTMV and wtTMV double-disk assembly shows an overlay of the backbones from the cross section of the assemblies. The inter-disk distance is shorter in dTMV (purple and grey monomers) compared with that in wtTMV (colored orange). Direct protein-protein interactions are present in the cavity region between disks of dTMV, in contrast to wtTMV which has no residue contacts in this region. (b) Superimposition of the backbones of a monomer from the flat disk (colored purple) and a monomer from the concave disk (colored grey) of dTMV only differs in the low-radius regions (red box). The concave monomers have a more ordered structure in that region.

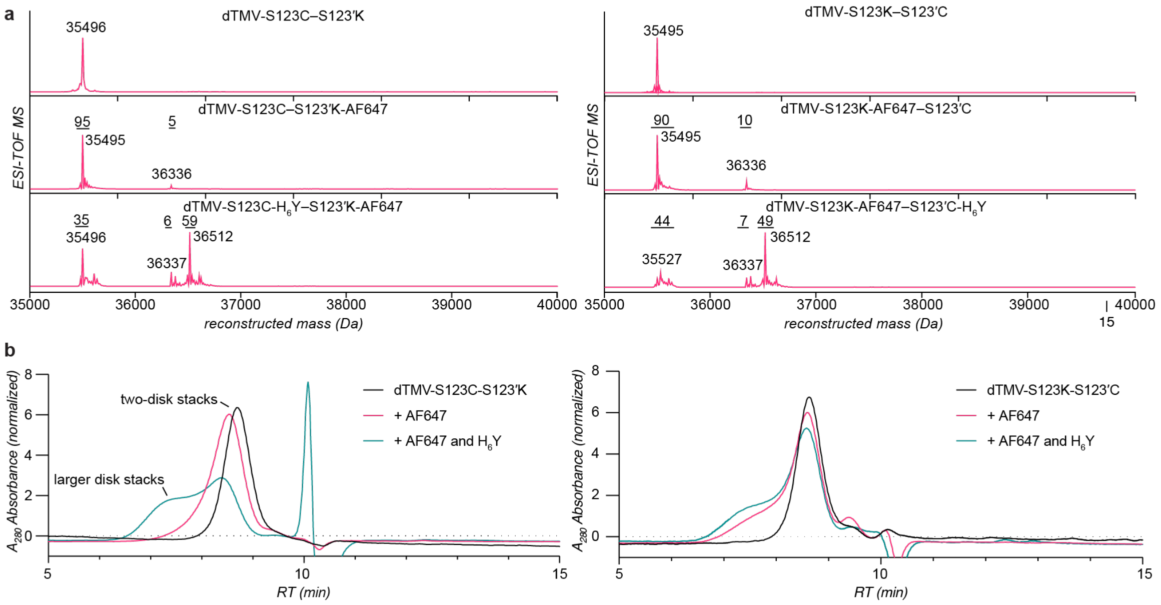


**Fig. S7.** Modification of dTMV with AlexaFluor 647 and $H_6Y$. (a) Mass data of dTMV-S123K–S123′C and dTMV-S123C–S123′K shows 5-10% modification with AlexaFluor 647, and 50-60% modification with the $H_6Y$ peptide. (b) Size exclusion chromatography of modified and unmodified dTMV shows a slight size increase upon modification with peptides, but little difference in assembly state between modified and unmodified proteins is indicated. Only fractions in the size range of two disk stacks were collected and used in subsequent experiments involving dTMV assembly association with SLBs.

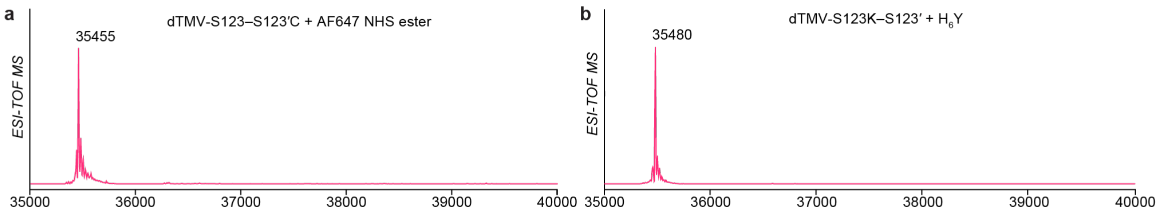


**Fig. S8.** Modification site of dTMV with AlexaFluor 647 and $H_6Y$. (a) Mass data of dTMV-S123–S123′C (MW: 35455) incubated with AlexaFluor 647 NHS ester (Expected MW: 36296) shows no modification without a lysine residue engineered at position S123. (b) Mass data of dTMV-S123K–S123′ (MW: 35481) incubated with the $H_6Y$ and tyrosinase (Expected MW: 36499) shows no modification without a cysteine residue engineered at position S123′.

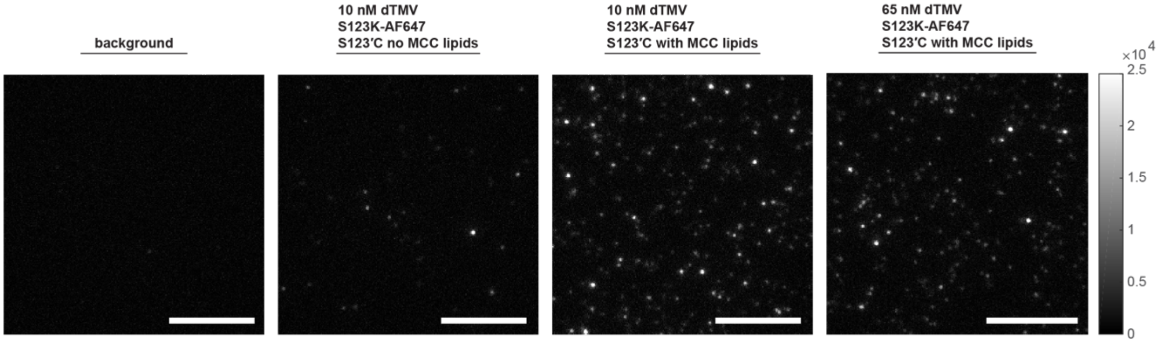


**Fig. S9.** Association of dTMV containing an engineered cysteine residue with SLBs. A minor amount of non-specific adherence to the SLB was observed when dTMV-S123K-AF647–S123′C was incubated with an SLB lacking lipids with maleimide head groups, the coupling partner. When dTMV-S123K-AF647–S123′C was incubated with maleimide-containing lipids at similar concentrations, the density of mobile complexes on the bilayer was not significantly higher than for the non-maleimide-containing SLBs. A higher density was also not achieved by raising the dTMV incubation concentration. Scale bars 10 µm.

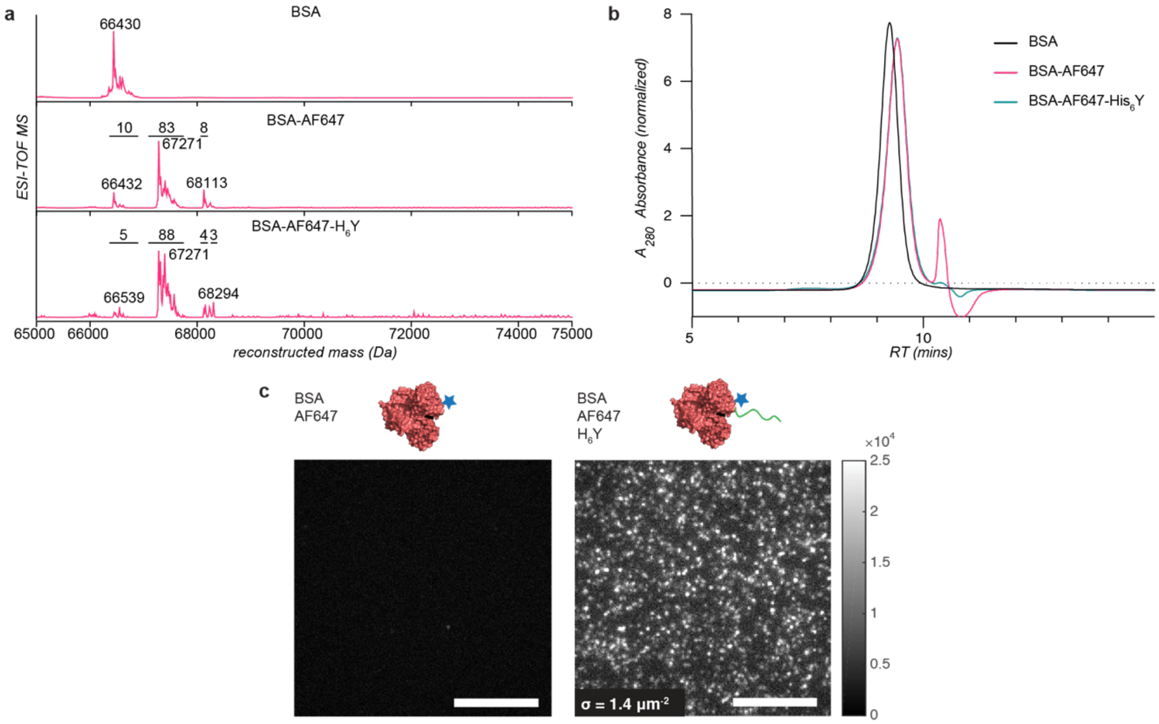


**Fig. S10.** Bovine serine albumin (BSA) conjugated to a His-tag and attached to supported lipid bilayers (SLBs). (a) Labeling of BSA with Alexafluor 647 (AF647) NHS ester resulted in 83% of proteins being labeled with a single dye and 8% being labeled with two dyes. The addition of the $H_6Y$ peptide to the AF647-labeled BSA resulted in approximately 3% of BSA monomers being labeled with a single $H_6Y$ peptide. (b) Size exclusion chromatography of unmodified and modified BSA shows a slight size decrease upon labeling with both dye and peptide. This may occur because BSA had dimerized pre-labeling, and labeling disrupted interactions at the homodimer interface. (c) Incubation of BSA on a supported lipid bilayer containing lipids with Ni-NTA headgroups resulted in sparse, nonspecific labeling in the sample without a His-tag and denser, specific bilayer labeling in the sample with BSA conjugated to the $H_6Y$ peptide, similar to the dTMV assembly results. Scale bars 10 μm.

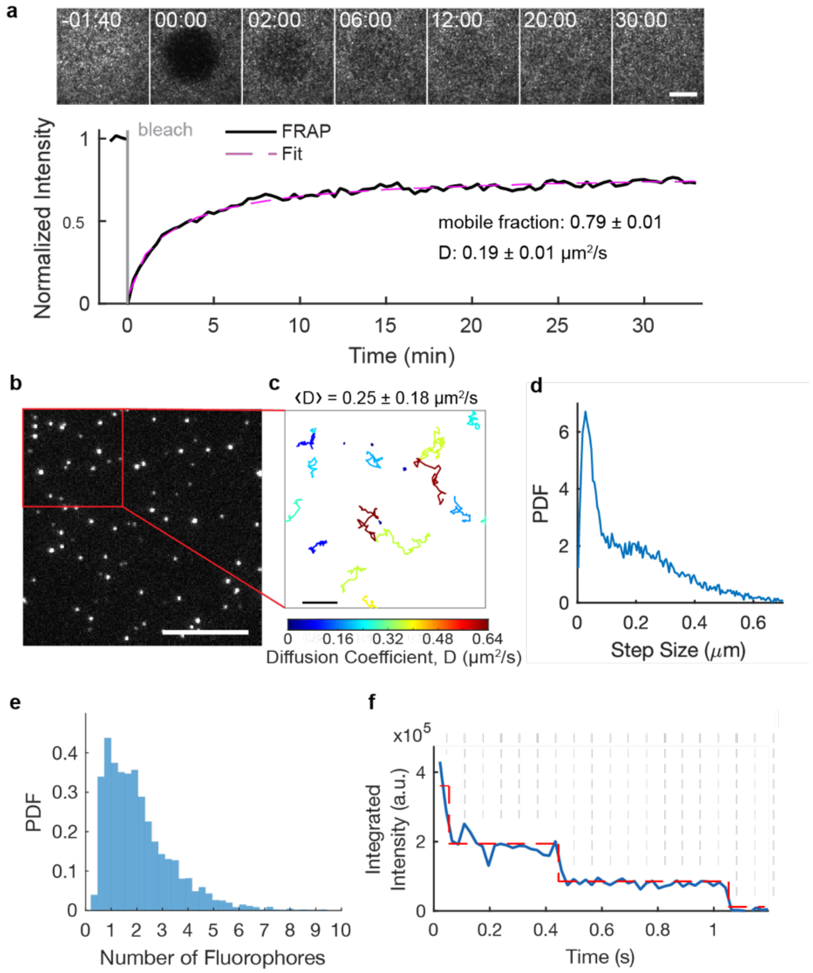


**Fig. S11.** dTMV S123K-AF647-S123′K-His6 is mobile on supported lipid bilayers. (a) Fluorescence recovery after photobleaching of a high density (13 $\mu m^{-2}$) of dTMV on the bilayer indicates that 79% of the dTMV disks on the bilayer are mobile, and that the mobile fraction diffuses at 0.19 ± 0.01 $\mu m^2/s$ on average. Scale bar 10 µm. Error denotes 95% CI. (b) A low density (0.2 $\mu m^{-2}$) of disks on the bilayer enables the analysis of single disk particles. Scale bar 5 µm. (c) Tracks of particles depicted in (b) diffusing through time illustrate the varied diffusion rates of particles on the bilayer. Tracks are colored by their diffusion coefficient, D. Scale bar 2 µm. Error denotes standard deviation. (d) The step size distribution of diffusing disks contains contributions from slowly-diffusing particles at smaller step sizes and quickly-diffusing particles at larger step sizes. n = 34,992 steps, n = 1,031 trajectories. (e) The background-subtracted fluorescence intensity distribution of all disks on the bilayer is calibrated to the mean integrated intensity of disks labeled with a single fluorophore. n = 3,338 particles. (f) A representative particle photobleaches in three steps, indicating that three fluorophores are conjugated to that particle. Scale bar 0.5 µm.

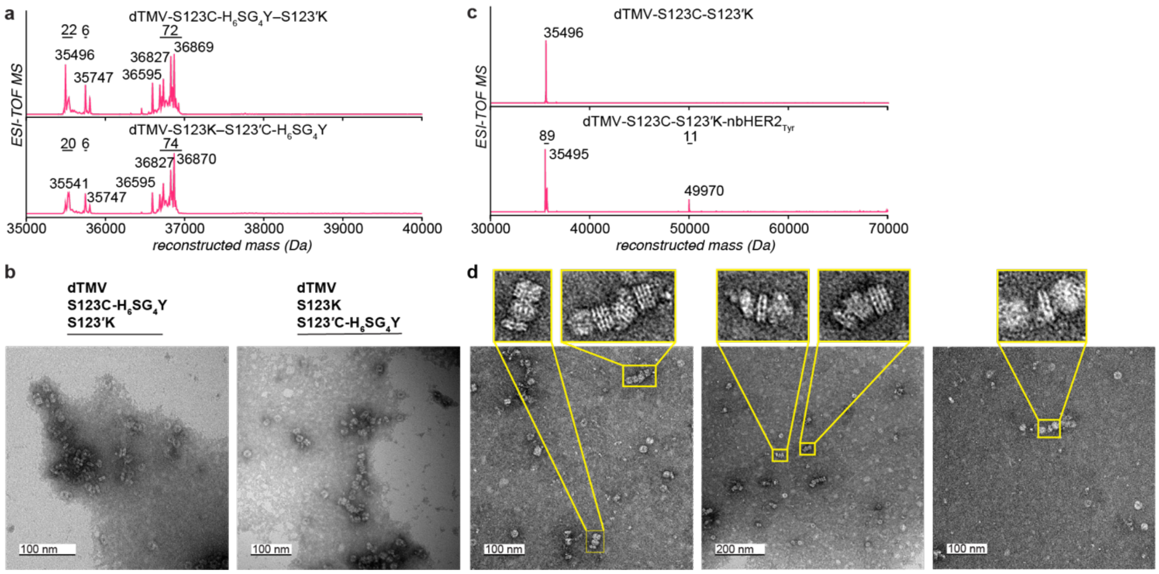


**Fig. S12.** TEM images of dTMV conjugated to an anti-HER2 nanobody and the $H_6SG_4Y$ peptide. Differences in attachment ability of mutants modified at nTMV versus the corresponding position on cTMV′ would contribute to understanding whether monomers are randomly oriented or uniformly oriented on the top and bottom disks. (a) Mass data of dTMV-S123C–S123′K and dTMV-S123K–S123′C conjugated to the peptide $H_6SG_4Y$ shows 70-75% of monomers were modified with peptide (expected MW of conjugate: 36829 Da). Additional peaks are likely due to peptide impurities. (b) Mass data of dTMV-S123C–S123′K conjugated to Human Epidermal Growth Factor Receptor 2 nanobody with a C-terminal SGGGGY tag ($nbHER2_{Tyr}$) shows that 11% of monomers were modified (expected MW of conjugate: 49973 Da), corresponding to an average of 1.9 copies of $nbHER2_{Tyr}$ per disk. (c) TEM images of dTMV-S123K–S123′C-$H_6SG_4Y$ and dTMV-S123C-$H_6SG_4Y$–S123′K look similar, both showing examples of interrupted stacks of disks. Few differences appear whether the $H_6SG_4Y$ peptide is attached to the Cys123 position on the nTMV or cTMV′ subunits. (d) TEM images of dTMV-S123C–S123′K conjugated via Cys123 to $nbHER2_{Tyr}$ show some examples of double disks with excess protein on either face.

**Table S1. X-ray crystallography data collection and refinement statistic.**

| | **dTMV**<br>(PDB code: 8EAW) |
|---|---|
| **Resolution range** | 78.84 - 2.8 (2.9 - 2.8) |
| **Space group** | C 2 2 21 |
| **Unit cell** | 208.51 255.5 260.85 90 90 90 |
| **Total reflections** | 1409281 (136305) |
| **Unique reflections** | 169799 (16816) |
| **Multiplicity** | 8.3 (8.1) |
| **Completeness (%)** | 99.74 (98.82) |
| **Mean I/sigma(I)** | 9.55 (1.18) |
| **Wilson B-factor** | 58.83 |
| **R-merge** | 0.2275 (1.877) |
| **R-meas** | 0.2422 (2.003) |
| **R-pim** | 0.08253 (0.6904) |
| **CC1/2** | 0.996 (0.467) |
| **CC*** | 0.999 (0.798) |
| **Reflections used in refinement** | 169652 (16702) |
| **Reflections used for R-free** | 8362 (788) |
| **R-work** | 0.1937 (0.3315) |
| **R-free** | 0.2274 (0.3637) |
| **CC(work)** | 0.955 (0.678) |
| **CC(free)** | 0.955 (0.593) |
| **Number of non-hydrogen atoms** | 38675 |
| **macromolecules** | 38658 |
| **solvent** | 17 |
| **Protein residues** | 4964 |
| **RMS (bonds)** | 0.004 |

| | |
|---|---|
| **RMS (angles)** | 0.65 |
| **Ramachandran favored (%)** | 97.29 |
| **Ramachandran allowed (%)** | 1.99 |
| **Ramachandran outliers (%)** | 0.72 |
| **Rotamer outliers (%)** | 0.07 |
| **Clashscore** | 3.42 |
| **Average B-factor** | 63.14 |
| **macromolecules** | 63.15 |
| **solvent** | 45.40 |

**Movie S1 (separate file).** Fluorescence recovery after photobleaching shows dTMV-S123C-$H_6$Y–S123′K-AF647 mobility on supported lipid bilayers. Images were taken at 20 s intervals to minimize photobleaching. Scale bar 10 μm.

**Movie S2 (separate file).** Fluorescence recovery after photobleaching shows dTMV- S123K-AF647–S123′C-$H_6$Y mobility on supported lipid bilayers. Images were taken at 20 s intervals to minimize photobleaching. Scale bar 10 μm.

**Movie S3 (separate file).** Single particles of dTMV-S123C-$H_6$Y–S123′K-AF647 diffuse within the two-dimensional supported lipid bilayer. Particles vary in brightness and mobility as a result of the non-stoichiometric labeling of AF647 and $H_6$Y to dTMV double disks. Particles tracks, corresponding to tracks shown in Fig. 5c, are shown in cyan. Scale bar 2 μm.

**Movie S4 (separate file).** Single particles of dTMV- S123K-AF647–S123′C-$H_6$Y diffuse within the two-dimensional supported lipid bilayer. Particles vary in brightness and mobility as a result of the non-stoichiometric labeling of AF647 and $H_6$Y to dTMV double disks. Particles tracks, corresponding to tracks shown in Fig. 5c, are shown in cyan. Scale bar 2 μm.

**SI References**

1. W. Kabsch, XDS. *Acta Crystallogr. Sect. D Biol. Crystallogr.* **66**, 125–132 (2010).
2. P. R. Evans, G. N. Murshudov, How good are my data and what is the resolution? *Acta Crystallogr. Sect. D Biol. Crystallogr.* **69**, 1204–1214 (2013).
3. P. A. Karplus, K. Diederichs, Linking Crystallographic Model and Data Quality. *Science (80-. ).* **336**, 1030–1033 (2012).
4. P. Emsley, B. Lohkamp, W. G. Scott, K. Cowtan, Features and development of Coot. *Acta Crystallogr. Sect. D Biol. Crystallogr.* **66**, 486–501 (2010).
5. P. V. Afonine, *et al.*, Towards automated crystallographic structure refinement with phenix.refine. *Acta Crystallogr. Sect. D Biol. Crystallogr.* **68**, 352–367 (2012).
6. D. Liebschner, *et al.*, Macromolecular structure determination using X-rays, neutrons and electrons: recent developments in Phenix. *Acta Crystallogr. Sect. D Struct. Biol.* **75**, 861–877 (2019).
7. M. Amblard, J.-A. Fehrentz, J. Martinez, G. Subra, Methods and Protocols of Modern Solid Phase Peptide Synthesis. *Mol. Biotechnol.* **33**, 239–254 (2006).
8. W.-C. Lin, C.-H. Yu, S. Triffo, J. T. Groves, Supported Membrane Formation, Characterization, Functionalization, and Patterning for Application in Biological Science and Technology. *Curr. Protoc. Chem. Biol* **2**, 235–269 (2010).
9. A. D. Edelstein, *et al.*, Advanced methods of microscope control using μManager software. *J. Biol. Methods* **1**, e10 (2014).
10. M. Carnell, A. Macmillan, R. Whan, “Fluorescence Recovery After Photobleaching (FRAP): Acquisition, Analysis, and Applications” in *Methods in Membrane Lipids*, Second Edi, D. M. Owen, Ed. (Humana Press, 2015), pp. 255–271.
11. J. J. Lin, *et al.*, Membrane Association Transforms an Inert Anti-TCRβ Fab’ Ligand into a Potent T Cell Receptor Agonist. *Biophys. J.* **118**, 2879–2893 (2020).
12. M. Axmann, G. J. Schütz, J. B. Huppa, Single Molecule Fluorescence Microscopy on Planar Supported Bilayers. *J. Vis. Exp.* (2015) https:/doi.org/10.3791/53158.
13. D. L. Ensign, V. S. Pande, Bayesian detection of intensity changes in single molecule and molecular dynamics trajectories. *J. Phys. Chem. B* (2010) https:/doi.org/10.1021/jp906786b.
14. M. T. Dedeo, K. E. Duderstadt, J. M. Berger, M. B. Francis, Nanoscale protein assemblies from a circular permutant of the tobacco mosaic virus. *Nano Lett.* **10**, 181–186 (2010).
15. B. Bhyravbhatla, S. J. Watowich, D. L. D. Caspar, Refined Atomic Model of the Four-Layer Aggregate of the Tobacco Mosaic Virus Coat Protein at 2.4-Å Resolution. *Biophys. J.* **74**, 604–615 (1998).
16. , Schrödinger Release 2022-1: Desmond Molecular Dynamics System (2021).

17. W. L. Jorgensen, J. Chandrasekhar, J. D. Madura, R. W. Impey, M. L. Klein, Comparison of simple potential functions for simulating liquid water. *J. Chem. Phys.* **79**, 926–935 (1983).
18. H. J. C. Berendsen, J. P. M. Postma, W. F. van Gunsteren, J. Hermans, "Interaction Models for Water in Relation to Protein Hydration BT - Intermolecular Forces: Proceedings of the Fourteenth Jerusalem Symposium on Quantum Chemistry and Biochemistry Held in Jerusalem, Israel, April 13–16, 1981" in B. Pullman, Ed. (Springer Netherlands, 1981), pp. 331–342.
19. C. Lu, *et al.*, OPLS4: Improving Force Field Accuracy on Challenging Regimes of Chemical Space. *J. Chem. Theory Comput.* **17**, 4291–4300 (2021).
20. G. J. Martyna, D. J. Tobias, M. L. Klein, Constant pressure molecular dynamics algorithms. *J. Chem. Phys.* **101**, 4177–4189 (1994).
21. G. J. Martyna, M. L. Klein, M. Tuckerman, Nosé–Hoover chains: The canonical ensemble via continuous dynamics. *J. Chem. Phys.* **97**, 2635–2643 (1992).
22. Schrödinger Release 2022-1: Bioluminate (2021).
